# Adipocyte-specific ablation of the Ca^2+^ pump SERCA2 impairs whole-body metabolic function and reveals the diverse metabolic flexibility of white and brown adipose tissue

**DOI:** 10.1101/2022.03.11.483886

**Authors:** Marco Bauzá-Thorbrügge, Elin Banke, Belén Chanclón, Eduard Peris, Yanling Wu, Saliha Musovic, Cecilia Jönsson, Peter Strålfors, Patrik Rorsman, Charlotta S. Olofsson, Ingrid Wernstedt Asterholm

**Author notes:** Corresponding authors Dr Charlotta Olofsson & Dr Ingrid Wernstedt Asterholm University of Gothenburg, The Sahlgrenska Academy, Dept. Neuroscience and Physiology, Section of Metabolic Physiology Postal adress: Box 432, SE-405 30 Göteborg, Sweden, (Ingrid Wernstedt Asterholm). Shared last authors (Charlotta S. Olofsson, Ingrid Wernstedt Asterholm). E-mail addresses.

## Abstract

**Objective:** Sarco/endoplasmic reticulum Ca^2+^-ATPase (SERCA) transports Ca^2+^ from the cytosol into the ER and is essential for appropriate regulation of intracellular Ca^2+^ homeostasis. The objective of this study was to test the hypothesis that SERCA pumps are involved in the regulation of white adipocyte hormone secretion and other aspects of adipose tissue function and that this control is disturbed in obesity-induced type-2 diabetes.

**Methods:** SERCA expression was measured in isolated human and mouse adipocytes as well as in whole mouse adipose tissue by Western blot and RT-qPCR. To test the significance of SERCA2 in adipocyte functionality and whole-body metabolism, we generated adipocyte-specific SERCA2 knockout mice. The mice were metabolically phenotyped by glucose tolerance and tracer studies, histological analyses, measurements of glucose-stimulated insulin release in isolated islets, and gene/protein expression analyses. We also tested the effect of pharmacological SERCA inhibition and genetic SERCA2 ablation in cultured adipocytes. Intracellular and mitochondrial Ca^2+^ levels were recorded with dual-wavelength ratio imaging and mitochondrial function was assessed by Seahorse technology.

**Results:** We demonstrate that SERCA2 is downregulated in white adipocytes from patients with obesity and type-2 diabetes as well as in adipocytes from diet-induced obese mice. SERCA2-ablated adipocytes display disturbed Ca^2+^ homeostasis associated with upregulated ER stress markers and impaired hormone release. These adipocyte alterations are linked to mild lipodystrophy, reduced adiponectin levels, and impaired glucose tolerance. Interestingly, adipocyte-specific SERCA2 ablation leads to increased glucose uptake in white adipose tissue while glucose uptake is reduced in brown adipose tissue. This dichotomous effect on glucose uptake is due to differently regulated mitochondrial function. In white adipocytes, SERCA2 deficiency triggers an adaptive increase in *Fgf21*, increased mitochondrial UCP1 levels, and increased oxygen consumption rate (OCR). In contrast, brown SERCA2 null adipocytes display reduced OCR despite increased mitochondrial content and UCP1 levels compared to wild type controls.

**Conclusions:** Our data suggest causal links between reduced white adipocyte SERCA2 levels, deranged adipocyte Ca^2+^ homeostasis, adipose tissue dysfunction and type-2 diabetes.

**Graphical abstract:** 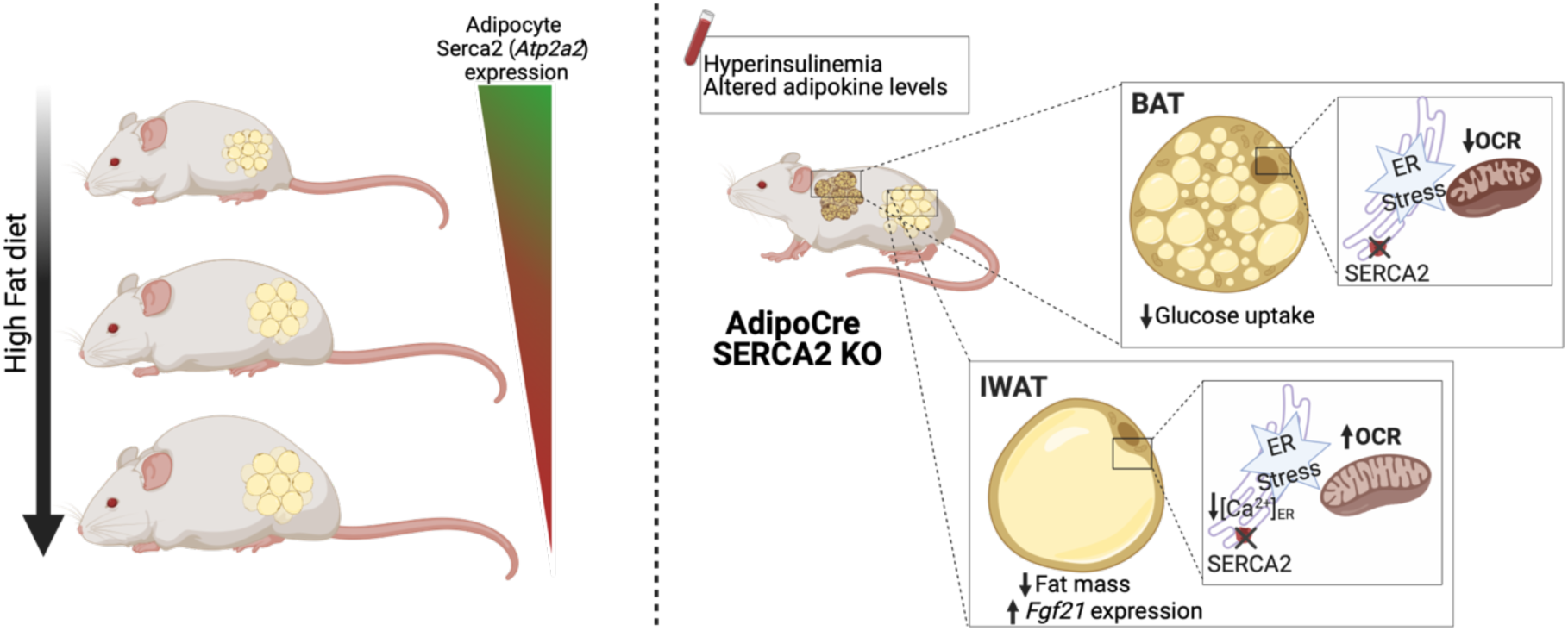

**Highlights:**

- Adipocyte SERCA2 is downregulated in human subjects with type-2 diabetes
- Loss of SERCA2 disturbs the intracellular Ca^2+^ homeostasis in adipocytes
- Impaired metabolism and altered adipokine levels in adipocyte-SERCA2 null mice
- Loss of SERCA2 accelerates metabolic processes in white adipocytes
- Loss of SERCA2 impairs the mitochondrial function of brown adipocytes

## 1. Introduction

The white adipose tissue (WAT) is important for maintanance of metabolic homeostasis in the body. Besides storing excess energy in the form of triglycerides, WAT also regulates whole-body metabolism by secreting a large variety of bioactive molecules, referred to as adipokines. Adipokines may be cytokines (e.g. interleukins and tumor necrosis factor alpha) whereas others, including adiponectin, leptin and resistin, are protein hormones. Obese adipose tissue is commonly dysfunctional with decreased triglyceride storage capacity and disturbed adipokine release, leading to ectopic lipid deposition, chronic low-grade inflammation and systemic insulin resistance [1; 2].

The endoplasmic reticulum (ER) constitutes the folding compartment for proteins entering a secretory pathway and is also a Ca^2+^ reservoir, important for controlling the dynamic regulation of Ca^2+^-dependent cellular functions including proliferation and differentiation, contraction, gene transcription, hormone secretion and apoptosis. Disrupted ER function causes mis- or unfolded proteins to accumulate within the ER and activates a process known as unfolded protein response (UPR). A high concentration of Ca^2+^ within the ER is essential for the normal function of chaperones and enzymes involved in protein folding and thus for the ability of UPR to resolve ER stress [3]. Moreover, intracellular Ca^2+^ has been shown to regulate several adipocyte-specific processes, such as lipolysis [4–6] and glucose uptake [7; 8]. Ca^2+^ is also important for regulation of the secretion of adipocyte hormones, such as the appetite-controlling hormone leptin [9–11]. In earlier studies, we defined an important regulatory role of Ca^2+^ for secretion of the adipocyte-specific insulin-sensitizing hormone adiponectin; Ca^2+^ potently augments cAMP-stimulated adiponectin exocytosis and is also required for sustained adiponectin release over prolonged time periods [12; 13].

The regulation of Ca^2+^ fluxes between ER and the cytosol is largely dependent on the sarco/endoplasmic reticulum Ca^2+^ ATPase (SERCA) pump that transports Ca^2+^ from the cytosol into the ER. Three paralogous genes code for SERCA1, 2 and 3. SERCA activity has also been linked to uptake of Ca^2+^ in the mitochondria, another important intracellular Ca^2+^ storage compartment [14]. The sequestration of Ca^2+^ within mitochondria impacts oxidative phosphorylation as well as cytosolic Ca^2+^ signals and excessive mitochondrial Ca^2+^ can trigger cell death (as reviewed in [15; 16]). Interestingly, studies in the heart, the liver and in pancreatic insulin-secreting beta-cells have affirmed SERCA pumps as key players in the pathophysiology of type-2 diabetes (T2D) [17–21]. However, diseases associated with disrupted ER Ca^2+^ dynamics have received insufficient attention and the role of SERCA for adipose tissue functionality is largely uninvestigated.

Here we show that SERCA2 is the paralog chiefly expressed in white mouse adipocytes isolated from both subcutaneous and visceral depots. Our study also demonstrates that SERCA2 is downregulated in adipocytes isolated from high fat diet (HFD)-induced obese mice and from human subjects with obesity and T2D. Moreover, genetic ablation of this Ca^2+^ pump results in a disturbed secretion of adipocyte hormones and impaired mitochondrial function in white and brown adipocytes, associated with whole-body glucose intolerance. Our study highlights an important causal link between a disturbed Ca^2+^ homeostasis in adipocytes and metabolic disease.

## 2. Material and Methods

### 2.1 Reagents and Tools Table

| REAGENT | SOURCE | IDENTIFIER |
| --- | --- | --- |
| **Antibodies** |  |  |
| Anti-Hsp60 antibody | Abcam | Ab46798 |
| Total OXPHOS Rodent WB Antibody Cocktail | Abcam | Ab110413 |
| Goat anti-mouse IgG | Abcam | Ab6788 |
| Anti-UCP1 antibody | Abcam | Ab10983 |
| Anti-SERCA2 ATPase antibody | Thermo fisher | MA3-919 |
| Anti- Beta-Tubulin | Cell signaling | 2128 |
| **Chemicals, Peptides, and Recombinant Proteins** |  |  |
| QIAzol | Qiagen | 79306 |
| Fast SYBR Green Master Mix | Thermo Scientific | 4385617 |
| (±)-Norepinephrine (+)-bitartrate salt (NE) | Sigma Aldrich | A0937 |
| Tamoxifen | Sigma Aldrich | T5648 |
| Bovine serum albumin | Sigma Aldrich | A7030 |
| DMEM, high glucose | Life Technology | 41965-039 |
| Seahorse XF DMEM Medium, pH 7.4 | Agilent | 103575-100 |
| Penicillin/streptomycin | Thermo Scientific | 15140-122 |
| Seahorse XF Glucose (1.0 M solution) / | Agilent | 103577-100 |
| Seahorse XF Pyruvate (100 mM solution) / | Agilent | 103578-100 |
| Seahorse XF L-Glutamine (200 mM solution) / | Agilent | 103579-100 |
| Dexamethasone | Sigma Aldrich | D4902 |
| Rosiglitazone | Sigma Aldrich | R2408 |
| Insulin Actrapid® Penfill® | Novo Nordisk | A10AB01 |
| 3-Isobutyl-1-methylxanthine (IBMX) | Sigma Aldrich | I5879 |
| DMSO | Sigma Aldrich | D5879 |
| PhosSTOP | Sigma Aldrich | 4906845001 |
| Hoechst | Thermo Scientific | H-21486 |
| CM-H2DCFDA | Thermo Scientific | C6827 |
| Mitotracker Green FM | Thermo Scientific | M7514 |
| Ultima Gold scintillation cocktail | Perkin Elmer | 6013326 |
| D-[U-<sup>14</sup>C]-glucose (5μCi/mouse, | Perkin Elmer | NEC042x25OU<br>C |
| Diisononyl phthalate | Sigma-Aldrich | **376663** |
| Fura-2 AM (Life Technologies) | Thermo Scientific | F1221 |
| Fetal Bovine Serum (FBS Hyclone) | Gibco | 10309433 |
| New Born Calf Serum (NBCS) | Gibco | 11530636 |
| RU360 | Sigma Aldrich | 557440 |
| Cyclopiazonic acid | Sigma Aldrich | C1530 |
| Forskolin | Sigma Aldrich | F6886 |
| CL 316243 disodium salt | Tocris | 1499 |
| ATP magnesium salt | Sigma Aldrich | A9187 |
| Thapsigargin | Tocris | 1138 |
| Eosin | Sigma Aldrich | HT110116 |
| Hematoxylin | Sigma Aldrich | GHS316 |
| *N,N,N',N'*-Tetramethyl-*p*-phenylenediamine (TMPD) | Sigma Aldrich | T7394 |
| Trans-Blot® Turbo™ Midi PVDF Transfer Packs | Bio-Rad | 1704157 |
| 10x Tris/glycine/sds | Bio-Rad | 161-0772 |
| L-Ascorbic acid | Sigma Aldrich | A0278 |
| Succinate | Sigma Aldrich | S2378 |
| Antimycin A | Sigma Aldrich | A8674 |
| Rotenone | Sigma Aldrich | R8875 |
| D-(+)-Malic acid | Sigma Aldrich | 02300 |
| pluriStrainer® (100µm) | Pluriselect | 43-50100-51 |
| Rhod-2, AM | Thermo Scientific | R1245MP |
| **Critical Commercial Assays** |  |  |
| BCA protein assay kit | Thermo Scientific | CAT#23225 |
| Seahorse XFe96 FluxPax | Agilent | CAT#102416-100 |
| High-Capacity cDNA Reverse Transcription Kit | Thermo Scientific | CAT#4368814 |
| Bradford | Amresco | M172-1L |
| Mouse Insulin ELISA kit | Mercodia | 10-1247-01, |
| Total adiponectin ELISA kit | EMD Millipore | EZMADP-60K |
| High molecular weight (HMW) adiponectin ELISA kit | MyBiosource | MBS028367 |
| Leptin DuoSet ELISA kit | R&D Systems | DY498 |
| Resistin DuoSet ELISA kit | R&D Systems | DY1069 |
| Total adiponectin DuoSet ELISA kit | R&D Systems | DY1119 |
| ReliaPrep™ RNA Cell Miniprep System | Promega | Z6112 |
| Seahorse XF Cell Mito Stress Test Kit | Agilent | 103015-100 |
| MouseWG-6 v2.0 Gene Expression BeadChips | Illumina | NA |
| **Experimental Models: Cell Lines** |  |  |
| 3T3L1 preadipocytes | Zenbio | SP-L1-F |
| **Experimental Models: Organisms/Strains** |  |  |
| loxP flanked SERCA2 knock-in *C57B6* mice | Geir Christensen (UiO, Norway) | N/A |
| Mouse Adipoq-Cre (B6; FVB-Tg(Adipoq-cre)1Evdr) | Jackson Laboratory | J stock 010803 |
| Mouse Adipoq-CreER (B6.129-Tg(Adipoq-cre/Esr1*)1Evdr) | Jackson Laboratory | J Stock 024671 |
| **Oligonucleotides** |  |  |
| RT qPCR Primers | This Paper | Table S1 |

| Software and Algorithms |  |  |
| --- | --- | --- |
| Graphpad Prism 7.0 for statistical analysis | GraphPad | N/A |
| Image Lab | Bio-Rad | N/A |

### 2.2 Experimental Model and Subject Details

#### 2.2.1 Genetically-Modified Mouse Lines

C57BL/6 background (Jackson Laboratory) loxP flanked SERCA2 knock-in mice were obtained from Geir Christensen (UiO, Oslo, Norway) and crossed with adiponectin promoter driven Cre transgenic mice with C57BL/6 background (B6; FVB-Tg (Adipoq-cre)1Evdr/J stock 010803, Jackson Laboratory) to selectively knock-out SERCA2 from adipocytes. Littermates not expressing Cre were used as wildtype controls. Male or female 8-9-week-old C57BL/6 mice (Jackson Laboratory) were fed chow (Global Diet #2016, Harlan-Teklad) or high fat diet (HFD, 60% kcal from fat; D12492, Research Diets Inc.) through 8 weeks. To obtain sufficient amounts of primary SERCA2 knockout adipocytes for *ex vivo* adipocyte hormone secretion experiments, SERCA2-loxP mice were crossed into CreER and adiponectin promoter-driven transgenic mice (Jackson Laboratory, B6.129-Tg (Adipoq-cre/Esr1*)1Evdr/J Stock 024671) to generate an inducible adipocyte-specific SERCA2 knockout mouse model. Tamoxifen (Sigma-Aldrich T5648) was dissolved in sunflower oil and given by gavage (5mg/mouse) at 10 weeks. After two weeks, the mice were euthanized and adipose tissue was dissected. All animal work was approved by the Regional Ethical Review Board in Gothenburg Sweden. All mice are male unless otherwise specified in the text, graph or legend.

#### 2.2.2 Mouse Adipocyte Cultures

3T3-L1 preadipocytes (ZenBio) cells were maintained as subconfluent cultures in DMEM (high-glucose, 4500 mg; Life Technologies, Stockholm, Sweden) containing 10% fetal bovine serum (Thermo Fisher Scientific, Waltham, MA, USA) and 1% penicillin–streptomycin (medium 1; Life Technologies) and differentiated into mature adipocytes as previously described [13]. Briefly, cells were grown to confluence (day 0) and thereafter incubated in medium 1 supplemented with 1 μM dexamethasone, 850 nM insulin, and 0.5 mM 3-isobutyl-1-methylxanthine (IBMX). After 48 h (day 2) the medium was changed to medium 1 supplemented with insulin only. After an additional 48 h (day 4) the medium was then replaced by medium 1 alone and was thereafter freshly replaced every 2 days. Experimental studies were performed in mature 3T3-L1 adipocytes between days 8 and 10 from the start of differentiation.

IWAT stromal vascular fraction (SVF) were isolated as previously described [22]. IWAT SVF were differentiated into mature adipocytes following the same protocol used for 3T3-L1 preadipocyte, but with the addition of 1 μM rosiglitazone (Sigma) in the first differentiation step. Brown fat SVF was isolated and differentiated as previously described [23]. The cell culture medium was DMEM, supplemented with 10% newborn calf serum (Thermo Fisher Scientific, Waltham, MA, USA), 4 nM insulin, 4 mM glutamine, antibiotics (50IU penicillin/mL and 50 μg streptomycin/mL), 10 mM HEPES, and 25 μg/mL sodium ascorbate (Sigma-Aldrich, St. Louis, MO, USA). The medium was changed on days 1, 3, 6 and 9. The SVF IWAT and BAT adipocytes cells were typically used for experiment at day 10 when they were fully differentiated as judged by their lipid content. For hypoxia treatment, adipocytes were cultured in 5% O2 in a InvivO2 300 hypoxia workstation (Baker Ruskinn, Bridgend, United Kingdom) at 37 °C. For normoxia controls, cells remained to be cultured in the humidified atmosphere (5% CO2, 95% air) at a temperature of 37 °C.

#### 2.2.3 Human Subjects

Informed consent was obtained from all participants after oral and written information. The procedures have been approved by the Regional Ethical Board, Linköping University, and were performed in accordance with the WMA Declaration of Helsinki. Adult women who were undergoing elective gynaecological abdominal surgery under general anaesthesia at the Department of Obstetrics and Gynaecology at the University Hospital in Linköping were recruited consecutively. A slice of abdominal subcutaneous adipose tissue from skin to muscle fascia was excised, and adipocytes were immediately isolated.

### 2.3 Experimental Procedures

#### 2.3.1 Isolation of human adipocytes and sample preparation

Human adipocytes were isolated from adipose tissue samples by collagenase (type 1, Worthington, NJ, USA) as described previously [24]. Adipocytes were lysed after separating cells from medium by centrifugation through dinonylphtalate oil. To minimize modification of proteins cells were immediately dissolved in SDS with protease and protein phosphatase inhibitors, frozen within 10 sec, and later thawed in boiling water for further processing for SDS-PAGE, as described [25].

#### 2.3.2 Oral glucose tolerance test

Glucose (2.5 mg/g mouse) was given by oral gavage after a 4 h fast and blood samples were collected at 0, 15-, 30-, 60- and 120-min. Adipose tissue and liver were dissected for further analysis with immunohistochemistry, RT-qPCR or western blotting as indicated.

#### 2.3.3 Glucose uptake

An oral load of D-[U-^14^C]-glucose (5μCi/mouse, Perkin Elmer, Boston, MA, USA) in 1.39M of glucose-PBS solution was given to adult mice by gavage after 4h fasting. After 2h, tissue was weighed and collected in a 2:1 chlorofom:methanol solution, homogenized in a tissue lyser and stored at 4°C overnight. CaCl_2_ 1M was added to all samples and centrifuged at 4°C 3000rpm during 20 min to separate aquous and organic phases. The incorporation of ^14^C was quantified in both phases using Ultima Gold scintillation cocktail (Perkin Elmer, Boston, MA, USA) and a beta counter (Perkin Elmer, Boston, MA, USA). Results are expressed as % of total ^14^C counts per mg tissue.

#### 2.3.4 Measurements of insulin and adipokines

Blood was collected and serum separated by clotting and centrifugation. Serum insulin was measured using Mouse Insulin ELISA kit (10-1247-01, Mercodia), total adiponectin by ELISA kit EZMADP-60K (EMD Millipore), high molecular weight (HMW) adiponectin by ELISA kit MBS028367 (MyBiosource), leptin by DuoSet ELISA DY498 and resistin by DuoSet ELISA DY1069 (R&D Systems). Adipokines were measured in tissue lysates obtained as previously described [26].

#### 2.3.5 Isolation of mouse adipose tissue mitochondria

Mitochondria of IWAT or BAT were isolated by differential centrifugation, based upon [27]. Freshly dissected adipose tissue was minced with scissors in ∼10 volumes of ice cold MSHE+BSA (70 mM sucrose, 210 mM mannitol, 5 mM HEPES, 1 mM EGTA and 0.5% (w/v) fatty acid-free BSA, pH 7.2, 4°C). The tissue was disrupted using a drill-driven Teflon glass homogenizer with 4–5 strokes. Homogenate was centrifuged at 800g for 10 min at 4°C, the lipid layer was carefully aspirated, and the remaining supernatant was separated and centrifuged at 8000 g for 10 min at 4°C. The pellet was resuspended in MSHE, and the centrifugation was repeated. The final pellet was resuspended for further analysis of oxygen consumption or western blot. Total protein was determined using Bradford Assay reagent (Bio-Rad).

#### 2.3.6 Pancreatic islets isolation

Pancreata were injected through the portal vein with 3-5 mL cold Hanks buffer containing 0.95 mg/mL collagenase V (Sigma, USA). The collected pancreata were incubated at 37°C for 7 min. The released islets were washed with Hanks buffer (0.1% BSA) 3-5 times and then hand-picked under a dissecting microscope to ensure that the islet preparation was pure. The islets were first cultured for 1 hour in RPMI 1640 containing 10% FBS, 100U/mL Penicillin/Streptomycin, 10 mM glucose. After washing the islets once with KRBH buffer + 0.1% BSA, 10-12 islets of similar size were handpicked in 0.3 mL KRBH buffer (0.1% BSA) containing 1 mM glucose. The islets were then pre-incubated KRBH buffer (0.1% BSA) containing 1 mM glucose for 30 min at 37°C, followed by 1-hour incubation in 0.3 mL KRBH buffer (0.1% BSA) containing (in mM): 1, 6 or 20 glucose, 20 glucose + 0.11 Tolbutamide or 20 glucose + 10 Arginine. Insulin was measured in the collected medium.

#### 2.3.7 Western blot analysis

10 μg protein from isolated mitochondria or 50 μg protein from isolated mouse adipocytes or equal volumes of packed human adipocytes were separated on SDS-PAGE gels and electrotransferred onto a PVDF membrane. The membranes were blocked with blocking solution (5% BSA in TBS, pH 7.5, containing 0.1% Tween 20) for 1 h and incubated overnight with the corresponding antibodies. Polyclonal antibodies against OXPHOS, UCP1, SERCA2, Beta-Tubulin, HSP60 were used as primary antibodies. Immunoblots were developed using an enhanced chemiluminescence kit. Bands were visualized with the ChemiDoc XRS system (Bio-Rad, CA, USA) and analysed with the image analysis program Image Lab© (Bio-Rad, CA, USA).

#### 2.3.8 Gene expression analysis

RNA was extracted and purified from adipose tissue and liver using QIAzol^®^ (Qiagen) and ReliaPrep^TM^ RNA Cell Miniprep System (Promega). RNA was reversely transcribed to cDNA by High-Capacity cDNA Reverse Transcription Kit. Fast SYBR Green Master Mix (Thermo Fisher Scientific, Waltham, MA, USA). 5 μg of the original RNA concentration and primers at a concentration of 500 nM were used in the qRT-PCR and gene expression was normalized against β-actin (*Atcb*) using the relative ΔCt method. Primer sequences are provided in **Suppl. Table 1.** Serca1-3 (*Atp2a1-3*) adipocyte and SVF mRNA expression data in **Suppl. Fig. 1A** was obtained by DNA microarray using MouseWG-6 v2.0 Gene Expression BeadChips (Illumina).

#### 2.3.9 Histology

Mouse adipose tissue, liver and pancreata were collected and fixed in 4% paraformaldehyde for 24 h. After paraffin embedding and sectioning (5μm for adipose tissue, liver and pancreas), tissues were stained with hematoxylin and eosin.

#### 2.3.10 [Ca^2+^]_i_ imaging

Differentiated 3T3-L1 were kept in an extracellular solution (EC) containing (in mmol/L): 140 NaCl, 3.6 KCl, 2 NaHCO_3_, 0.5 NaH_2_PO_4_, 0.5 MgSO_4_, 5 HEPES (pH 7.4 with NaOH), 2.6 CaCl_2_, and 5 glucose. Cells were loaded with Fura-2 AM (Life Technologies) and [Ca^2+^]_i_ values were recorded with dual-wavelength ratio imaging as previously described [28]. Excitation wavelengths were 340 and 380 nm, and emitted light was collected at 510 nm. The absolute [Ca^2+^]_i_ was calculated using Eq. 5 in the study by Grynkiewicz et al (K_d_= 224 nmol/L) [29]. For measurements of [Ca^2+^] inside mitochondria ([Ca2+ ]mit), cells were incubated with 5 μM rhod-2/AM and 0.02% pluronic F-127 for 30 min at 37 ◦C [30]. Rhod-2/AM was excited at 543 nm and emission was recorded at 576 nm.

#### 2.3.11 Adiponectin and resistin secretion in primary adipocytes

Adiponectin and resistin secretion was measured in mouse primary adipocytes isolated from inguinal white adipose tissue (30-min incubations at 32°C). Primary adipocytes was isolated from 8-week-old mice inguinal white adipose tissue. Tissue samples were minced and digested using collagenase type II (1 mg/mL; 45–60 min; 37°C) and poured through a 100-µm nylon mesh. Floating adipocytes were washed with Krebs-Ringer glucose buffer (1% BSA) and immediately used for experiments. Primary adipocytes were diluted to 25% volume/volume. Secretion was measured in an EC (see [Ca^2+^]_i_ imaging) containing test substances as indicated. Primary adipocytes cells were separated from media by centrifugation in diisononyl phthalate (Sigma-Aldrich) followed by snap freezing in dry ice. Tubes were cut through the oil layer at two points. Cells were lysed in PBS containing SDS (2%) and protease inhibitor (1 tablet/10mL; PhosSTOP, Sigma Aldrich). EC aliquots and cell homogenates were stored at −80°C.

Secreted adiponectin or resistin (measured with mouse ELISA DuoSets; R&D Systems) was expressed in relation to total protein content (BCA protein assay).

#### 2.3.12 Oxygen consumption rate

For the Mitostress test, Agilent instructions were followed. Briefly, adipocytes differentiated onto XF96 Microplates (Seahorse Bioscience, Agilent Technologies, CA, USA) from IWAT or BAT SVF were washed with assay medium and incubated for 1 hour at 37°C. The oxygen consumption rate (OCR) was determined at four levels: with no additions, and after adding oligomycin (2 μM), carbonyl cyanide 4- (trifluoromethoxy) phenylhydrazone (FCCP, 2 μM), and antimycin A + rotenone (0.5 μM both). Basal oxygen consumption rate and mitochondrial Oxidative Phosphorylation System (OXPHOS) parameters were measured according to manufacturer’s instructions of the Mito Stress test Kit (Seahorse Bioscience, Agilent Technologies, CA, USA). Results were normalized to protein quantified using a bicinchoninic acid-based protein assay (Thermo Fisher Scientific, Waltham, MA, USA) in each well.

For the Electron Flow Assay, Agilent instructions were followed. Briefly 5 µg of isolated mitochondria (BAT or IWAT) were resuspended in assay buffer (MAS-1; 70 mM sucrose, 220 mM mannitol, 2 mM HEPES, 1 mM EGTA, 10 mM KH_2_PO_4_, 5 mM MgCl_2_ and 0.2% BSA, pH 7.2), plus glutamate and malate (5 μM both). The mitochondria were seeded, centrifuged and incubated 30 minutes at 37°C. After basal readings, 2.2 μM Rotenone was injected to inhibit electron transport complex I. In subsequent additions, succinate (5 mM) was added as a substrate of electron transport complex II followed by 40 μM Antimycin A to inhibit complex III. Finally, 5 mM Ascorbate and 10 mM N,N,N0,N0-Tetramethyl-p-phenylenediamine (TMPD) were added to measure the Cytochrome *c* oxidase activity (complex IV).

For the coupling between the electrons, 5 µg of isolated IWAT mitochondria were resuspended in MAS-1 buffer, seeded, centrifuged and incubated 30 min at 37°C. To examine de degree of coupling between the electron transport chain and the oxidative phosphorylation machinery, coupled state was measured with the presence of substrate succinate (5 mM) and 2 μM rotenone (State II). State III was initiated with de addition of ADP (0.2 mM), meanwhile state IV was induced with the addition of oligomycin (1 μM). Maximal uncoupler-stimulated respiration was induced with the addition of 2 μM FCCP (State III). Finally, 0.5 μM antimycin A was added to block all mitochondrial respiration. The respiratory control ratio was calculated following the formula (RCR: State III/State IV).

For the norepinephrine (NE) OCR stimulation, differentiated BAT SVF brown adipocytes were incubated 1 hour at 37°C, in assay media with no BSA, previous to start of the readings. After basal measurements, NE (1 μM) was injected in order to measure adrenergically induced uncoupled respiration. NE-induced uncoupled respiration curves were calculated by subtracting the basal OCR from the OCR values after NE treatment.

#### 2.3.13 Measurements of mitochondrial content and intracellular ROS

Imaging experiments were carried out on a Zeiss LSM 710 confocal microscope Mitotracker Green (Thermo Fisher Scientific, Waltham, MA, USA) was used at 200nM for 90min, meanwhile Hoechst (Thermo Fisher Scientific, Waltham, MA, USA) was used at 2μg/μl for 10 min followed by wash-out before imaging. Mitotracker Green was excited by a 488 nm laser, while 361 nm laser was used for detection of Hoechst.

Imaging was performed with a 40× objective. The dye signal was quantified with the Image J analysis software and normalized to the number of cells (Hoechst dye). The ROS assay was performed as previously described [31]. Values were expressed as the ratio between the fluorescence of the oxidized CM-H2DCFDA (Thermofisher) and Hoechst, after subtraction of the signal of not-stained cells.

### 2.4 Statistical Analysis

Statistical analysis was performed with GraphPad Prism 8.0 (GraphPad Software, CA, USA). Statistical parameters, including n values, are noted in figure legends. Data are presented as means ± standard error of the mean (SEM) and a p < 0.05 was considered significant. Statistical analysis of data with more than 2 groups was assessed with one-way ANOVA. Data with two groups was analyzed by two-tailed Student’s *t* test.

## 3. Results

### 3.1 White adipocyte SERCA2 is down-regulated in HFD-induced obese mice and in obese type-2 diabetic humans

By DNA microarray analysis, we found that *Serca2* (*Atp2a2*) and -3 (*Atp2a3*) are the most abundant *Serca*-paralogs in mouse inguinal, gonadal and mesenteric WAT (IWAT, GWAT and MWAT) with expression both in the adipocyte and in the stromal vascular fraction (SVF), while *Serca1* (*Atp2a1*) was detected at extremely low levels. In isolated adipocytes, *Serca2* was the predominant paralog, although *Serca3* was also highly expressed (**Suppl. Fig. 1A**). SERCA2 expression and/or activity are reduced in liver, pancreatic islets and cardiomyocytes in animal models of obesity and T2D [18; 32; 33]. This prompted us to investigate whether HFD-induced obesity reduces *Serca2* also in adipose tissue. Indeed, in both IWAT and GWAT *Serca2* mRNA levels were reduced from the fourth week in HFD-induced obese mice. In contrast, *Serca3* mRNA levels were unaltered in both fat depots until week 16 of HFD, when the expression was dramatically decreased in IWAT but not in GWAT (**Fig. 1A-D**). This HFD-induced decrease in WAT *Serca2* was due to reduced levels specifically in adipocytes, while SVF levels of *Serca2* remained unaffected (**Fig. 1E-F**). The expression of *Serca*3 was unaltered in the isolated IWAT adipocyte fraction and tended to be downregulated in SVF from this adipose tissue depot (**Fig. 1G**). *Serca*3 mRNA levels were however upregulated in GWAT adipocytes and unchanged in the SVF fraction (**Fig. 1H**). In contrast to findings in WAT, the brown adipose tissue (BAT) *Serca2* expression was slightly elevated in HFD-induced obese mice while *Serca*3 expression was unaltered (**Fig. 1I**). To determine whether adipocyte SERCA2 is relevant also for human pathophysiology, we measured SERCA2 protein in subcutaneous adipocytes obtained from subjects with and without obesity-associated T2D. SERCA2 levels were ∼20% lower in adipocyte from the obese individuals compared to non-obese controls (**Fig. 1J**) and a correlated inversely with plasma glucose levels of adipocyte donors (**Fig. 1K).** As hypoxia downregulates SERCA2 in cardiomyocytes [34–36], we hypothesized that a similar mechanism may be at play in enlarged hypoxic adipocytes of insulin-resistant subjects [37; 38]. Indeed, hypoxia led to reduced *Serca2* expression in 3T3-L1 adipocytes (**Fig. IL**). In conclusion, our results pinpoint *Serca2* as the SERCA paralog that is downregulated in white adipocytes from both diet-induced obese mice and obese/T2D patients.

**Figure 1.**
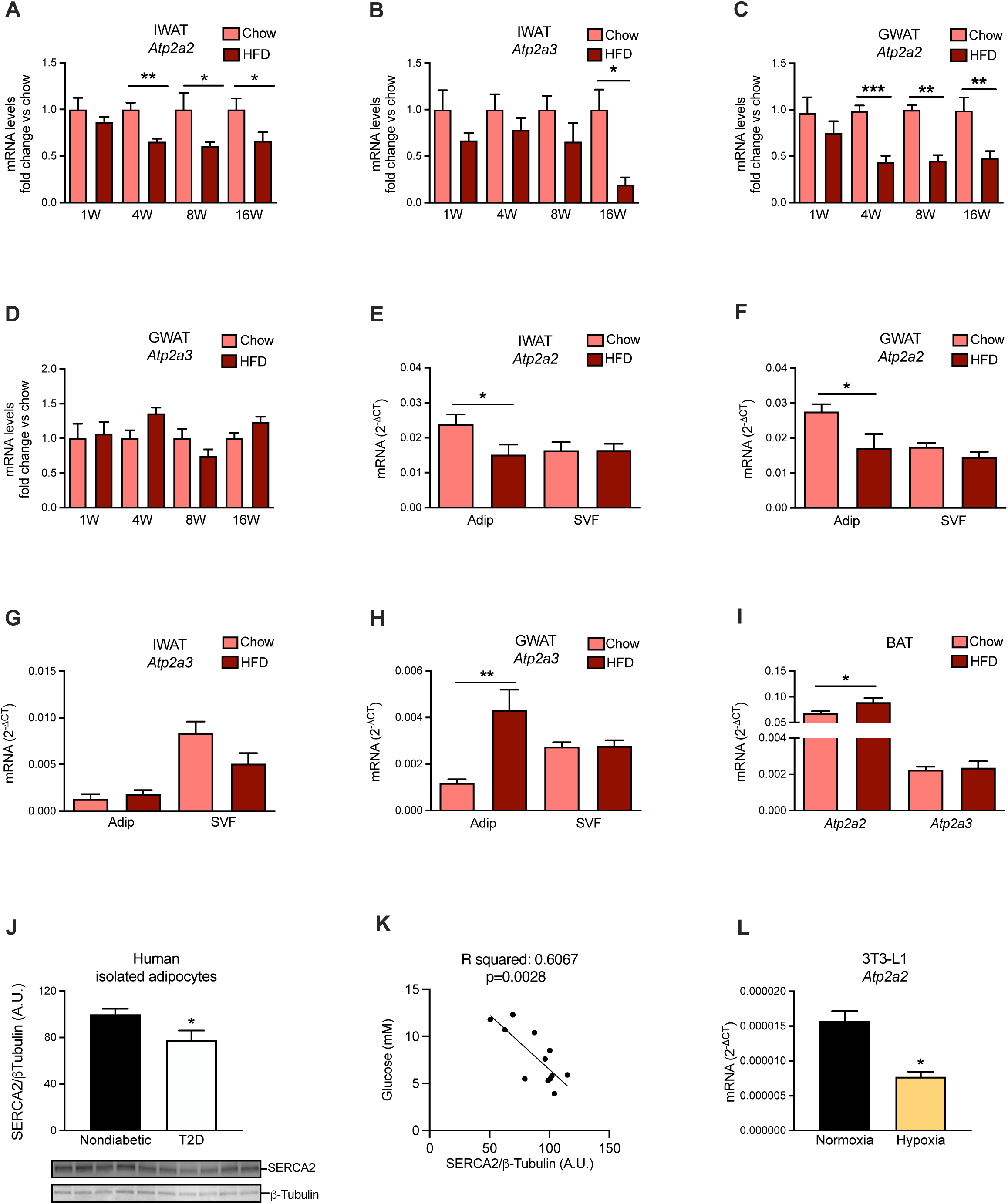
Adipocyte SERCA2 is down-regulated in high fat diet-induced obesity. *Serca2 (Atp2a2)* and *3 (Atp2a3)* mRNA levels in (**A-B**) IWAT and (**C-D**) GWAT at 1, 4, 8 and 16 weeks (W) high fat diet (HFD)- and chow-fed wild type mice (N=6-10). (**E-H**) *Serca2 (Atp2a2) and 3 (Atp2a3)* mRNA levels in 10-week-HFD-fed mice IWAT and GWAT adipocytes or SVF (N=6-10). (**I**) *Serca2 (Atp2a2)* and *3 (Atp2a3)* gene expression in 10-week-HFD-fed mice BAT (N=4-6). (**J**) SERCA2 protein levels in human adipocytes isolated from patients diagnosed with type-2 diabetes (N=6) [mean age 67 years (range 65–72); mean BMI 36.9 kg/m2 (range 30–43)] and from nondiabetic subjects (N= 6) [mean age 64 years (range 55–69); mean BMI 24.3 kg/m2 (range 23–27)]. (**K**) Correlation analysis for donor plasma glucose with SERCA2 in primary human adipocytes isolated from corresponding patients. (**L**) *Serca2 (Atp2a2)* gene expression in 3T3-L1 adipocytes at normoxia vs. 24h hypoxia (N=6/group). All values are expressed as mean ± SEM * p<0.05, **p<0.01, ***p<0.001 for HFD vs. chow.

### 3.2 Inhibition of SERCA in adipocytes is associated with altered Ca^2+^ homeostasis and defective adiponectin exocytosis

We next tested how SERCA inhibition affects the adipocyte Ca^2+^ homeostasis by performing ratiometric time-lapse recordings of intracellular Ca^2+^ concentrations. Pre-treatment of 3T3-L1 adipocytes with 20 μM of the SERCA inhibitor cyclopiazonic acid (CPA) reduced the ER calcium store by ∼50%, as shown by the diminished Ca^2+^ increase in response to acute addition of thapsigargin (another SERCA inhibitor) during the recording (**Fig. 2A-B**). A similar reduction of stored ER calcium was found in cultured adipocytes genetically ablated for SERCA2 (**Fig. 2C-E)**. The reduced Ca^2+^ stores in SERCA2-ablated adipocytes were associated with the transcriptional upregulation in IWAT of two key players of cellular Ca^2+^ influx mediated via tore- operated Ca^2+^ entry (SOCE): the stromal interaction molecule 1 (*Stim1*) that recognized when ER Ca^2+^ levels are low, and the calcium release-activated calcium channel protein 1, *Orai1* [39] (**Suppl. Fig. 2A**). We also observed a downregulation of the Ryanodine Receptor channel (*Ryr3*) that mediate Ca^2+^ release from ER to cytosol [40] (**Suppl. Fig. 2A**).

**Figure 2.**
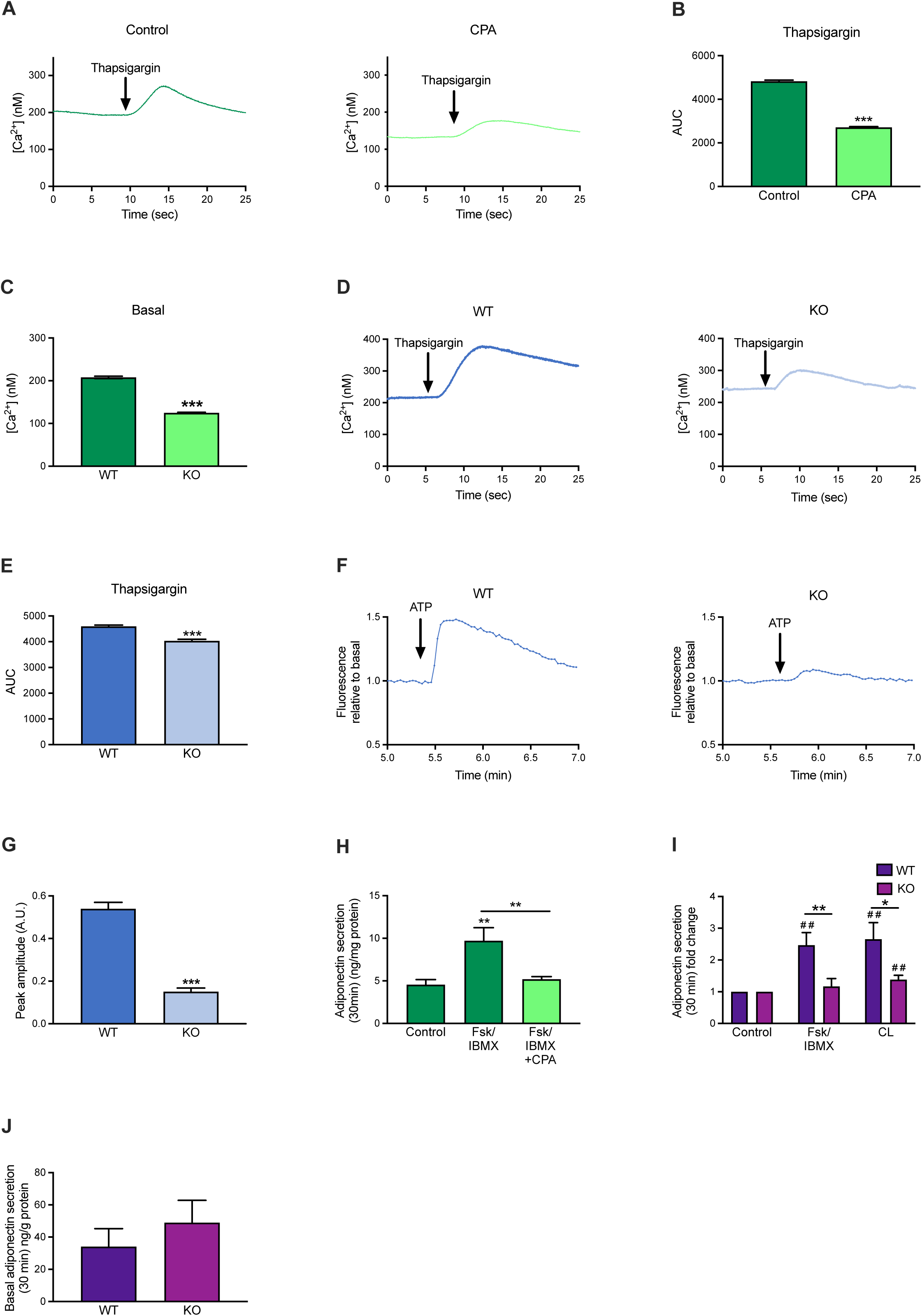
Inhibition of SERCA in adipocytes is associated with altered Ca^2+^ homeostasis and defective adiponectin exocytosis. (**A**) Example traces of typical [Ca^2+^]_i_ responses in 3T3-L1 using Fura-2 and exposed to slow flow of thapsigargin (1 µM) previously treated with cyclopiazonic acid (CPA) or vehicle for 4h. (**B**) Area under the curve (AUC) of the [Ca^2+^] response in 3T3-L1 using Fura-2 and exposed to slow flow of thapsigargin (1µM) previously treated with CPA or vehicle for 4h. (**C**) Basal [Ca^2+^] in 3T3-L1 using Fura-2 treated with CPA or vehicle for 4h. (**D**) Example traces of typical [Ca^2+^]_i_ responses in differentiated wildtype (WT) or SERCA2 knockout (KO) IWAT SVF cells using Fura-2 and exposed to slow flow of thapsigargin (1µM). (**E**) Area under the curve (AUC) of the [Ca^2+^] response in differentiated WT or adipocyte-specific SERCA2 KO IWAT SVF cells using Fura-2 and exposed to slow flow of thapsigargin (1µM). (**F**) Example traces in response to ATP of cytoplasmic Ca^2+^ in differentiated WT or SERCA2KO IWAT SVF cells using Fluo-4 dye. (**G**) ATP peak amplitude of the cytoplasmic Ca^2+^ in differentiated IWAT SVF cells using Fluo-4 dye. (**H**) Total adiponectin secretion during 30 min treatment with Forskolin (10µM; Fsk/IBMX) together with IBMX (200µM) in 3T3-L1 with CPA or vehicle for 24h. (**I**) Stimulated and (**J**) basal adiponectin secretion (30min) in primary IWAT adipocytes isolated from adipocyte-specific SERCA2 KO mice. Cells were incubated with (10µM; Fsk/IBMX) together with IBMX (200µM) or beta-3 adrenergic receptor agonist CL316,243 (10µM, CL). Values (N=6-10) are expressed as mean ± SEM *p<0.10, **p<0.05, ***p<0.01 for WT vs. KO.

It has been demonstrated that the extracellular application of ATP rapidly elevates cytosolic Ca^2+^ in adipocytes via purinergic P2Y2 receptors and STIM1/ORAI1-mediated activation of SOCE [41; 42]. We thus tested the effect of ATP on cultured SERCA2-ablated adipocytes. Mitochondria can accumulate a 10-fold higher concentration of Ca^2+^ than that measured in the cytosol and Ca^2+^ is transferred between ER and the mitochondria via mitochondria-associated membranes (MAMs), specialised regions of communication between the two organelles [43]. Mitochondrial Ca^2+^ levels were therefore measured in parallell. In agreement with [41; 42], extracellular addition of ATP (100 μM) elevated [Ca^2+^]_i_ in control adipocytes while the ATP-induced [Ca^2+^]_i_ peak was reduced in SERCA2-deficient adipocytes (**Fig. 2F-G**). The ATP-triggered mitochondrial Ca^2+^ elevations were similarly affected in these knockout adipocytes (**Suppl. Fig. 2B-C**). This finding was associated with a transcriptional upregulation of the mitochondrial calcium uniporter (*Mcu*) that mediates electrophoretic Ca²⁺ uptake into the mitochondrial matrix [44] in IWAT of adipocyte-specific SERCA2 deficient mice (**Suppl. Fig. 2A**). Thus, SERCA2 regulates Ca^2+^ in both mitochondria and the ER of white adipocytes.

To further explore the importance of SERCA for metabolic regulation, we studied the role of these Ca^2+^ pumps in adiponectin secretion. Ca^2+^ is important for sustained adiponectin secretion [12; 45] and both mitochondrial dysfunction and ER stress have been linked to reduced adiponectin production in adipocytes [46; 47]. The effect of SERCA inhibition on regulated adiponectin exocytosis has however not previously been studied. CPA treatment induced ER stress in 3T3-L1 adipocytes (as verified by increased *Bip* and *Xbp1s* expression), reaching significance after 4 hours (**Suppl. Fig. 2D-E)**. Adiponectin secretion stimulated by a combination of forskolin and IBMX (elevates intracellular cAMP and also slightly increases Ca^2+^ in adipocytes; [13]) was blunted in CPA-treated cells (**Fig. 2H)** whereas basal (unstimulated) adiponectin release was unaffected (not shown). Adiponectin secretion triggered by forskolin/IBMX or the beta 3 adrenergic receptor agonist CL-316,243 (CL) was similarly blunted also in primary adipocytes genetically ablated for SERCA2 (**Fig. 2I)**. Basal adiponectin release was not altered in SERCA2 null adipocytes compared to wild type (**Fig. 2J**).

### 3.3 Adipocyte-specific SERCA2 knockout causes ER stress in IWAT and BAT

To define the role of SERCA2 for adipocyte functionality and whole-body metabolic health, we generated an adipocyte-specific *Serca2* knockout mouse model. The knockouts displayed ∼60-70% reduced *Serca2* levels in IWAT, GWAT and BAT, while the SVF *Serca2* expression was unaffected (**Fig. 3A**). SERCA2 protein levels were reduced to a similar degree in adipocytes from different depots in SERCA2 knockout mice (**Fig. 3B**). There was a compensatory upregulation of *Serca3* expression in the adipocyte fraction of WAT and BAT, and this *Serca3* increase was particularly pronounced in GWAT adipocytes (**Fig. 3C**). Downregulation of *Serca2* was associated with increased expression of the UPR-markers *Xbp1s* and *Bip* and the inflammatory markers *Emr1* (*F4/80*), *Ccl2* (*Mcp1*) and *Tnfα* in IWAT and BAT, indicating ER stress (**Fig. 3D-E**). In support of a higher degree of IWAT inflammation, there was also an increased abundance of crown-like structures (**Fig. 3F**). Interestingly, the GWAT expression of UPR and inflammation markers was unaffected by knockdown of *Serca2* (**Suppl. Fig. 3**), possibly due to the compensatory elevation of *Serca3* in this fat depot (**Fig. 3C**).

**Figure 3.**
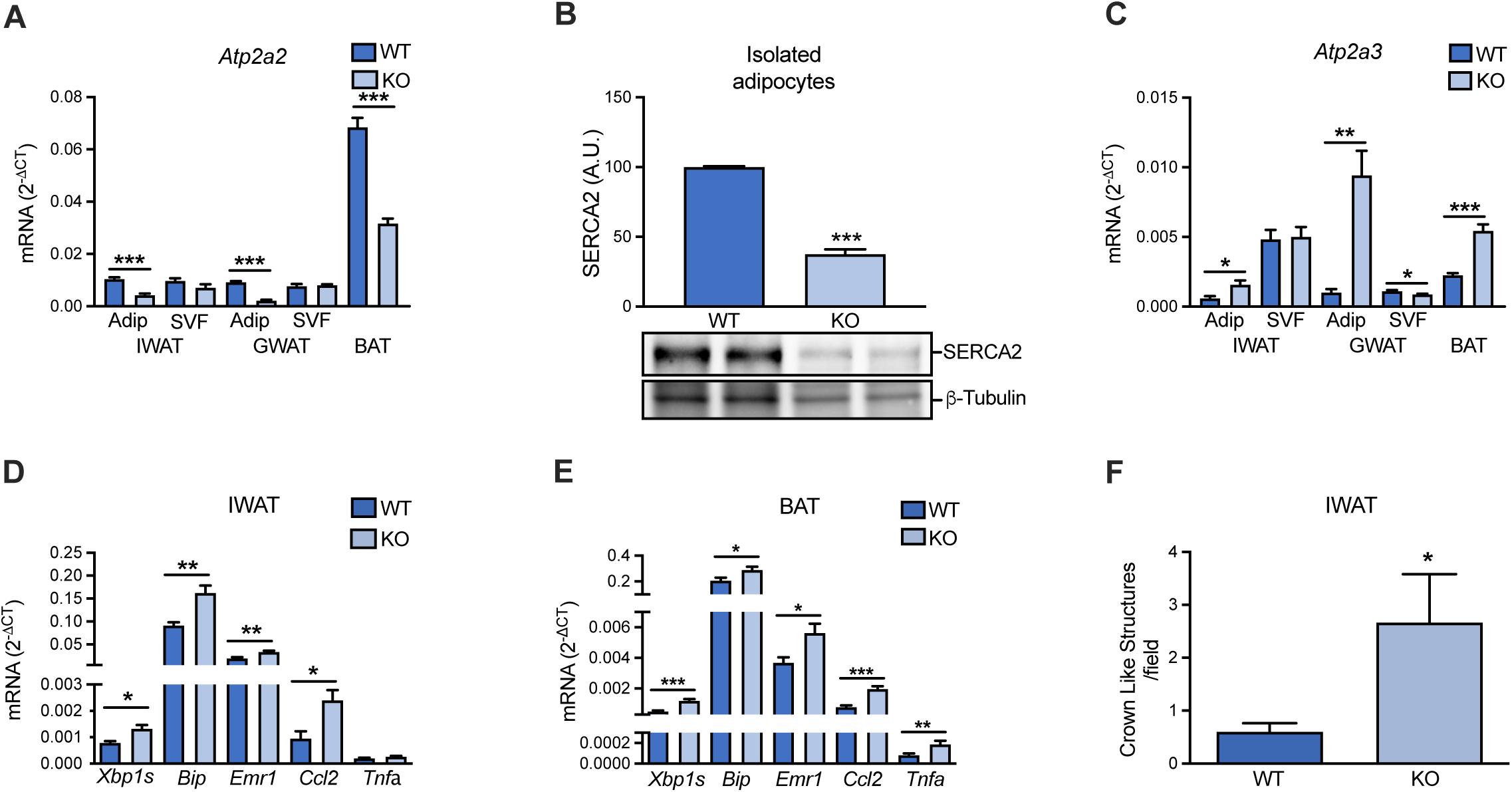
Adipocyte-specific SERCA2 ablation leads to elevated markers of ER stress and inflammation in IWAT and BAT. (**A**) *Serca2 (Atp2a2)* gene expression in adipocytes and SVF of IWAT, GWAT and BAT in chow diet-fed mice.**(B)** SERCA2 protein levels in isolated WT and adipocyte-specific SERCA2 KO IWAT adipocytes. (**C**) *Serca3 (Atp2a3)* gene expression in adipocytes and SVF of IWAT, GWAT and BAT in chow diet-fed mice. (**D-E**) ER stress and inflammation gene expression markers in IWAT and BAT chow diet-fed mice. (**F**) Crown like structures in IWAT of chow or HFD-fed mice. Values (N=6-10) are expressed as mean ± SEM *p<0.10, **p<0.05, ***p<0.01 for WT vs KO.

### 3.4 Adipocyte-specific SERCA2-deficient mice are mildly lipodystrophic and display whole-body metabolic dysfunction

Male and female adipocyte-specific SERCA2 knockout mice grew normally after weaning, but displayed reduced WAT mass, while the liver, pancreas and BAT mass was increased compared to littermate controls (**Fig. 4A-C)**. This lipodystrophic phenotype was associated with impaired whole-body metabolic function, as judged by slightly reduced glucose tolerance and hyperinsulinemia (**Fig. 4D-G)**, as well as a trend towards increased glucagon levels (2.61 ± 0.33 vs. 4.44 ± 0.97 pmol/L, p=0.122). The enlarged pancreata of adipocyte-specific SERCA2 knockout mice displayed normal morphology (judged by haematoxylin and eosin staining), and the islet density and size distribution per analyzed section were similar between genotypes (**Suppl. Fig 4A-C**). Insulin secretion induced by high glucose, arginine and tolbutamide was elevated in adipocyte-specific SERCA2 knockouts (**Suppl. Fig. 4D**). The increased BAT weight appeared to primarily result from increased fat deposition, as judged by the whiteish BAT appearance (**Fig. 4H)**. The hepatic triglyceride content was similar between genotypes (**Fig. 4I-J**).

**Figure 4.**
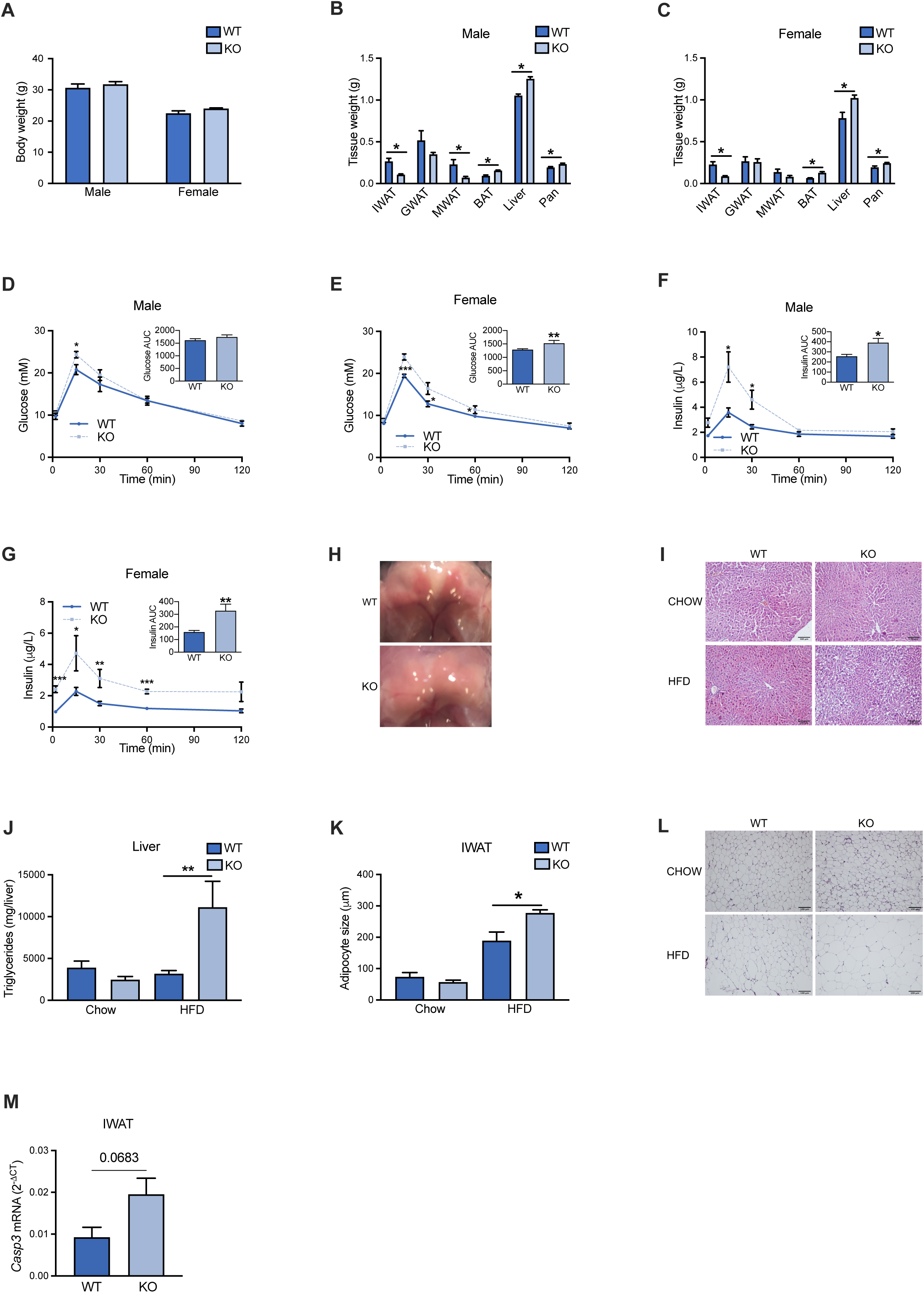
Adipocyte-specific SERCA2 deficient mice are lipodystropic and display impaired whole-body metabolic function. (**A**) Body weight, (**B-C**) IWAT, GWAT, MWAT, BAT, liver and pancreas weight of WT and adipocyte-specific SERCA2 KO chow-fed male and female mice at 8 weeks of age (N=6-10). (**D-G**) Circulating glucose and insulin levels at indicated time points and AUC in response to oral glucose load in chow-fed male and female WT and adipocyte-specific SERCA2 KO mice. (**H**) Representative images of BAT from WT and adipocyte-specific SERCA2 KO mice. (**I**-**J**) Liver triglyceride content and (**K-L**) adipocyte size in IWAT from mice on chow or 10-week-HFD. (**M**) *Casp3* in IWAT adipocytes from chow diet-fed mice. Values (N=6-10) are expressed as mean ± SEM *p<0.05, **p<0.01, ***p<0.001 for WT vs KO.

When fed HFD (to induce obesity), control and SERCA2 knockout mice exhibited similar weight gain (**Suppl. Fig 4E**), and the difference in WAT, BAT and liver mass persisted between genotypes (**Suppl. Fig 4F)**. Hepatic lipid deposition was increased in adipocyte-specific SERCA2 knockout mice (**Fig. 4I-J**). The glucose intolerance was aggravated by HFD and the differences between genotypes were similar to those observed at chow conditions (**Suppl. Fig. 4G-J**).

To elucidate the underlying reasons for the reduced WAT mass, we measured IWAT adipocyte size. During chow conditions, the adipocyte-specific SERCA2 knockouts displayed smaller IWAT adipocytes (although the difference did not reach significance). Despite ∼60% decreased IWAT weight, the SERCA2 knockout adipocytes of HFD-fed mice were ∼50% larger than littermate wild type control adipocytes (**Fig. 4K-L)**. This increase in average adipocyte size was primarily due to a doubling of the percentage of larger adipocytes (>350 μm in diameter), while the fraction of smaller adipocytes was lower (**Suppl. Fig. 4K**). These data indicate that SERCA2 deficiency does not lead to impaired lipid storage capacity of an individual adipocyte, but that there is an increase in HFD-induced adipocyte death (as implied by the increased occurrence of crown-like structures, **Fig. 3F**) and/or a decreased formation of new adipocytes in these mice. Given that only adiponectin-expressing cells (i.e. mature adipocytes), are SERCA2-deficient in this mouse model and that HFD-induced expansion largely depends on hypertrophy [48], we argued that increased adipocyte death is more likely to explain the fewer but larger IWAT adipocytes in HFD-fed mice. Indeed, the adipogenic capacity was similar between genotypes as judged by *ex vivo* differentiation of IWAT SVF cells (**Suppl. Fig. 4L)** and there was a tendency towards increased *Casp3* expression in IWAT of adipocyte-specific SERCA2 knockout mice (**Fig. 4M**)

### 3.5 Levels and secretion of adipocyte hormones are altered in adipocyte-specific SERCA2 knockout mice

As SERCA2-deficient adipocytes display abrogated regulated release of adiponectin (**Fig. 2H-I)**, we determined whether adipocyte-specific SERCA2 knockouts have altered levels of the adipose tissue hormones adiponectin, resistin and leptin. Both male and female adipocyte-specific SERCA2 knockout mice had much reduced levels of circulating adiponectin and this was associated with decreased adiponectin protein and gene expression in IWAT and GWAT (**Fig. 5A-B**, **Suppl. Fig. 5A)**. In contrast, serum HMW adiponectin levels were similar between adipocyte-specific SERCA2 knockouts and littermate wild type controls. However, HFD-feeding reduced both total and HMW adiponectin in both genotypes (**Fig. 5C**). As a consequence of the unaltered HMW adiponectin levels, the HMW to total adiponectin ratio was significantly higher in the male knockouts with a similar trend in females (**Suppl. Fig. 5B**). The ER chaperones disulfide-bond A oxidoreductase-like protein (*DsbA-L*), ER membrane-associated oxidoreductase (*Ero1-Lα*) and its associated protein *Erp44* have been identified as critical players in the assembly and secretion of HMW adiponectin [49; 50]. We found increased gene expression of *Ero1lα* in IWAT from both male and female knockouts, and higher gene expression of *Erp44* in IWAT from the female SERCA2 null mice (**Suppl. Fig. 5C**).

**Figure 5.**
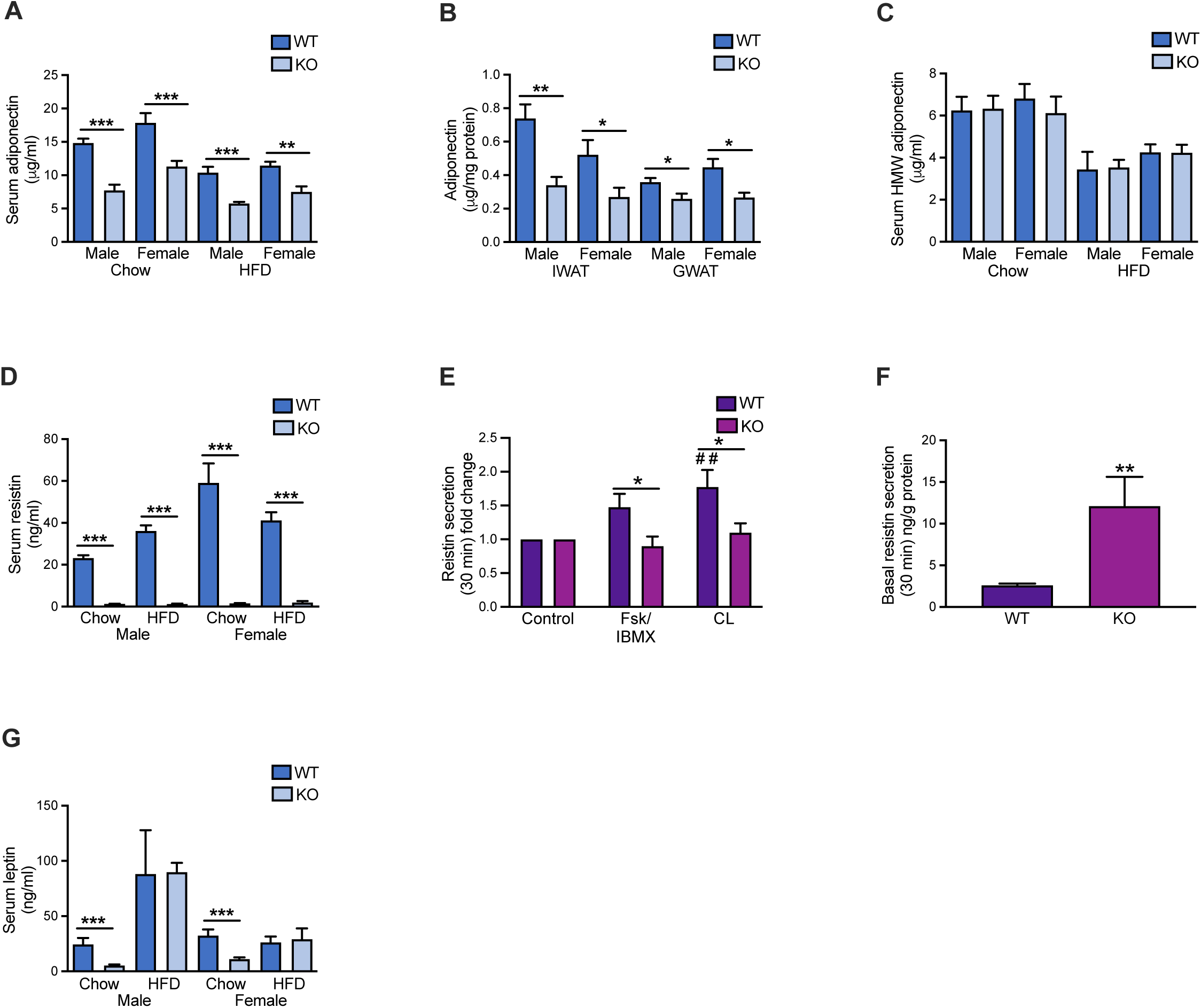
Adipocyte hormones are altered in adipocyte-specific SERCA2 knockout mice. (**A**) Serum total adiponectin levels, (**B**) IWAT and GWAT total adiponectin levels, (**C**) serum HMW-adiponectin levels, (**D**) serum resistin levels in male and female chow and 10-week-HFD-fed WT and adipocyte-specific SERCA2 KO mice. (**E**) Stimulated and (**F**) basal Resistin secretion (30min) in primary IWAT adipocytes isolated from adipocyte-specfic SERCA2 KO knockout. Cells were incubated with (10µM; Fsk/IBMX) together with IBMX (200µM) or beta-3 adrenergic receptor agonist CL316,243 (10µM, CL). (**G**) Serum leptin in male and female chow and 10- week-HFD-fed WT and adipocyte-specific SERCA2 KO mice. All values (N=6-10) are expressed as mean ± SEM *p<0.05, **p<0.01, ***p<0.001 for WT vs KO.

Resistin has been suggested to contribute to the pathogenesis of metabolic diseases, although its exact role is not fully elucidated [51–53]. Similar to adiponectin, resistin has a complex structure with multimeric assembly [54] and the two hormones are in mouse adipocytes co-released from the same vesicles [55]. As expected, circulating resistin levels were dramatically reduced both at chow and HFD-fed conditions (**Fig. 5D**) and the stimulated resistin secretion was blunted in primary SERCA2-ablated adipocytes, while the basal release was increased (**Fig. 5E-F**); IWAT and GWAT Resistin (*Retn*) mRNA levels were also much reduced (**Suppl. Fig. 5D**).

In contrast to adiponectin and resistin, leptin is released as a monomer in proportion to the degree of adiposity. We therefore expected a less potent effect of SERCA2 ablation on leptin levels. In line with this assumption, circulating leptin levels as well as IWAT and GWAT leptin (*Lep*) mRNA levels were similar between genotypes in HFD-fed mice (**Fig. 5G and Suppl. Fig. 5E**). Circulating leptin and IWAT *Lep* mRNA levels were however reduced in chow-fed knockouts, in agreement with their lower fat mass (**Fig. 5G and Suppl. Fig. 5E**).

### 3.6 Opposite effect of adipocyte-specific SERCA2 ablation on white and brown adipose tissue glucose uptake

We speculated that the phenotype of SERCA2 knockout mice is due to the systemic impact of impaired metabolic function of white and brown adipocytes. To investigate this, we used [^14^C]-labeled glucose to measure tissue glucose uptake and *de novo* lipogenesis 2h after an oral glucose load. Contrary to our expectations, we found that the glucose uptake was increased in WAT (represented by IWAT, GWAT and MWAT), while only BAT displayed a clear reduction in glucose uptake (**Fig. 6A**). The enhanced glucose uptake in knockout WAT was primarily due to increased [^14^C]-counts in the aqueous phase, likely reflecting glucose oxidation (**Fig. 6A**). In contrast, [^14^C]-counts in the organic phase (representing *de novo* lipogenesis) dominated in BAT (**Fig. 6A**). Glucose uptake was also augmented in liver, heart, skeletal muscle and pancreas, largely due to an increase in the aqueous phase (**Fig. 6A**). Thus, the impaired glucose uptake in BAT appears to be compensated for by increased glucose-stimulated insulin release from pancreas leading to enhanced glucose uptake in insulin-sensitive tissues.

**Figure 6.**
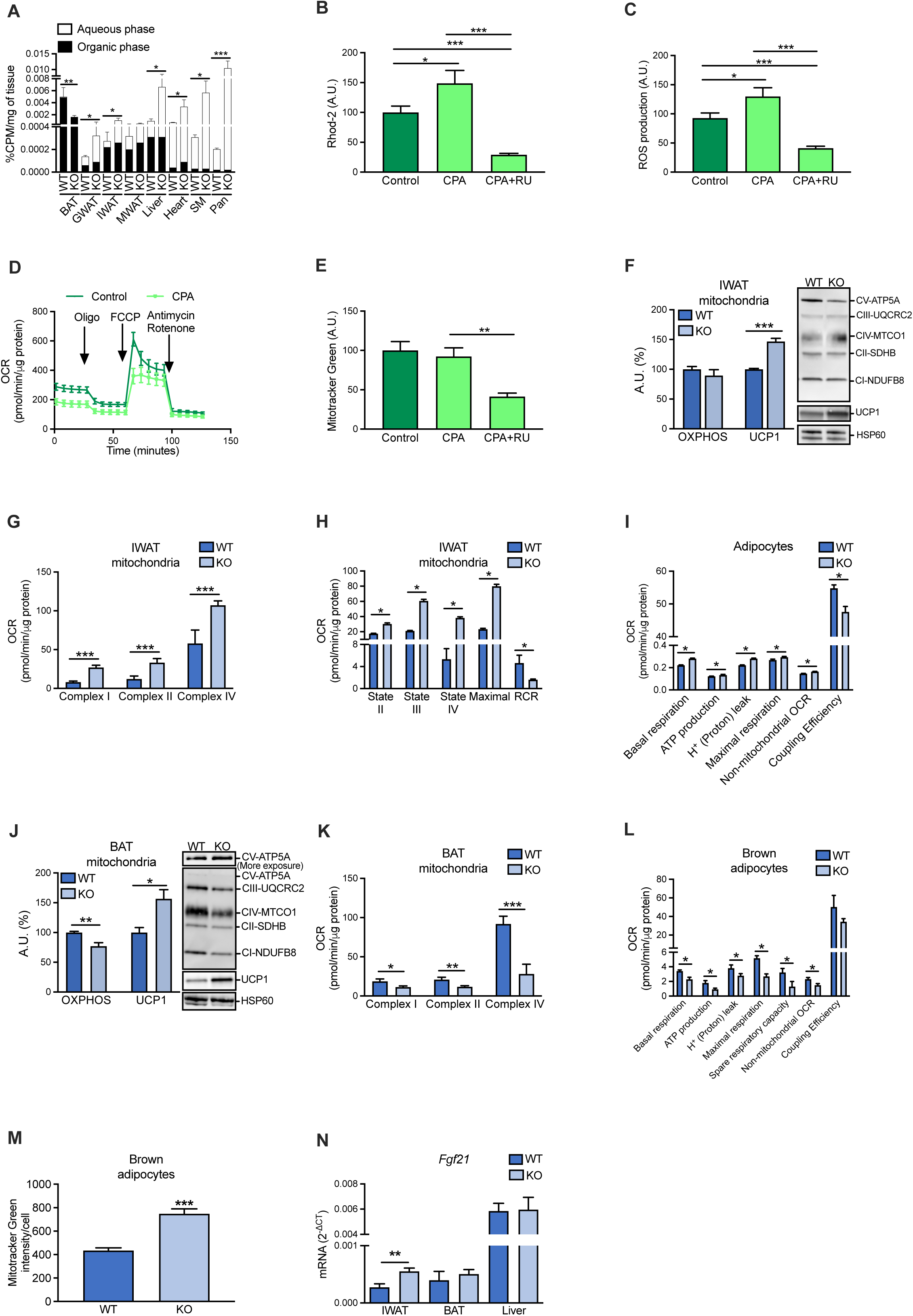
Opposite effect of adipocyte-specific SERCA2 ablation on white and brown adipose tissue glucose uptake and mitochondrial function. (A) Total glucose uptake in chow-fed WT and adipocyte-specific SERCA2 KO mice. (B) Mitochondrial calcium measured with Rhod-2 dye and (C) ROS production measured with CM-H_2_DCF-DA dye in 3T3-L1 treated with vehicle or CPA with or without the mitochondrial calcium uptake inhibitor RU for 1h. (D) Oxygen consumption rate (OCR) in 3T3-L1 treated with CPA or vehicle for 4h. (E) Mitochondrial mass measured with Mitotracker green dye. (F) Total OXPHOS (Complex I-V) and UCP1 protein levels in isolated mitochondria IWAT in chow-fed WT and adipocyte-specific SERCA2 KO mice. (G) OCR of the different mitochondrial complexes and (H) respiratory states analysed in isolated IWAT mitochondria from chow-fed WT and adipocyte-speecific SERCA2 KO mice. (I) OCR parameters calculated in adipocytes differentiated from IWAT SVF from chow-fed WT and adipocyte-speecific SERCA2 KO mice. (J) Total OXPHOS (Complex I-V) and UCP1 protein levels in isolated BAT mitochondria from chow-fed WT and adipocyte-speecific SERCA2 KO mice. (K) OCR of the different mitochondrial complexes in isolated BAT mitochondria from chow-fed WT and adipocyte-specific SERCA2 KO mice. (L) OCR parameters and (M) mitochondrial mass, measured with Mitotracker green dye, in brown adipocytes differentiated from BAT SVF from chow-fed WT and adipocyte-speecific SERCA2 KO mice. (N) *Fgf21* expression in IWAT, BAT and liver in chow-fed WT and adipocyte-specific SERCA2 KO mice. All values (N=3-6) are expressed as mean ± SEM *p<0.05, **p<0.01, ***p<0.001 vs WT.

### 3.7 Pharmacological inhibition of SERCA is associated with mitochondrial dysfunction in 3T3-L1 adipocytes

The increased glucose uptake and metabolism in WAT from adipocyte-specific SERCA2 null mice urged us to investigate the effect of SERCA inhibition on mitochondrial function. Imaging of Rhod-2 loaded 3T3-L1 adipocytes showed that the basal mitochondrial Ca^2+^ accumulation was increased by CPA treatment (**Fig. 6B**), indicating transfer of Ca^2+^ from the ER to the mitochondria [43]. The increase in mitochondrial Ca^2+^ was accompanied by a higher ROS production and reduced oxygen consumption rate (OCR; **Fig. 6C-D**). As expected, administration of the mitochondrial Ca^2+^ uptake inhibitor RU360 reduced both basal and CPA-induced mitochondrial Ca^2+^ and ROS levels (**Fig. 6B-C**). As shown in **Fig. 6E**, the cell mitochondrial content (judged by Mitotracker green staining) was unchanged by CPA, but reduced by RU360 treatment. Taken together, our results indicate that acutely disturbed intracellular Ca^2+^ homeostasis affects the mitochondrial function in adipocytes.

### 3.8 Adipocyte-specific SERCA2-deficient mice display altered mitochondrial dysfunction in IWAT and in BAT

Lack of SERCA2 in white adipocytes may result in enhanced metabolism (as indicated by the increased WAT glucose uptake) through a mitohormesis-like process. We explored this possibility by analyzing the expression of browning markers and found a trend for increased IWAT *Ucp1* levels in the adipocyte-specific SERCA2 knockout mice, while the browning markers *Prdm16* and *Dio2* were downregulated (**Suppl. Fig. 6A**). Notably, UCP1 protein levels in isolated IWAT mitochondria were ∼1.5-fold higher in SERCA2 ablated mice than in littermate wild type controls, while electron transport chain protein levels were similar between genotypes (**Fig. 6F**). Furthermore, the OCR of isolated IWAT mitochondria was much higher (**Fig. 6G-H**). This increase in mitochondrial OCR was primarily due to a lower coupling efficiency in SERCA2- deficient adipocytes (**Fig. 6H**). To test if this effect of SERCA2 ablation on mitochondrial function is cell autonomous, we analyzed the mitochondrial function in *in vitro* differentiated adipocytes from IWAT SVF. Indeed, also cultured SERCA2- deficient adipocytes displayed elevated uncoupled OCR and the ATP production-linked OCR was slightly increased (**Fig. 6I**).

The altered expression of browning markers was similar in BAT and IWAT; BAT *Prdm16* and *Dio2* levels were reduced, while *Ucp1* was unchanged in adipocyte-specific SERCA2 knockouts compared to littermate wild type controls (**Suppl. Fig. 6B**). Moreover, the level of total electron transport proteins was reduced along with increased UCP1 protein levels in isolated mitochondria from BAT of adipocyte-specific SERCA2 knockout mice (**Fig. 6J).** In sharp contrast to SERCA2- deficient white adipocytes, mitochondria of BAT adipocyte-specific SERCA2 knockout mice displayed lower OCR (**Fig. 6K**). Accordingly, OCR was decreased in cultured SERCA2-deficient brown adipocytes, as shown by the reduction of basal and ATP-production-linked respiration and other key mitochondrial respiration parameters, **(Fig. 6L)** despite a higher mitochondrial content (**Fig. 6M).** However, the capacity to increase the respiration in response to norepinephrine was unaffected (**Suppl. Fig. 6C**). Several studies have shown that ER stress can lead to an adaptive increase in fibroblast growth factor 21 (FGF21) [56; 57], a metabolic hormone capable of inducing thermogenesis in BAT and promoting a brown fat-like thermogenic program in white adipocytes [58; 59]. In line with the increased UPR marker expression (**Fig. 3D**), we found that *Fgf21* mRNA levels were elevated in adipocyte-specific SERCA2 knockout IWAT (**Fig. 6N**). However, such an ER-stress/UPR-associated induction of *Fgf21* was not observed in BAT (**Fig. 3E**, **Fig. 6N**). As expected, the hepatic *Fgf21* mRNA levels were similar between genotypes (**Fig. 6N**). Thus, our data suggest that genetic ablation of SERCA2 leads to different adaptations in white and brown adipocytes.

## 4. Discussion

Here we show that diet-induced obesity in mice, T2D in humans and hypoxia are associated with lower SERCA2 levels in white adipocytes. SERCA2-deficient white adipocytes display altered calcium homeostasis, hormone release and mitochondrial function, and adipocyte-specific SERCA2 knockout mice present with hyperinsulinemia, glucose intolerance and lower adiponectin levels, likely driven by both white and brown adipocyte dysfunction. In brief, the WAT of this mouse model fails to expand properly, presumably because of increased adipocyte death. However, the remaining SERCA2-deficient white adipocytes show accelerated metabolism, as judged by increased glucose uptake and increased OCR. In contrast, BAT is enlarged but less metabolically active, as demonstrated by its whiter appearance, decreased glucose uptake and decreased OCR. Furthermore, this mouse model illustrates how genetically normal tissues to some extent can compensate for adipose tissue defects.

For instance, the glucose uptake is dramatically increased in liver, heart, skeletal muscle and pancreas, and the glucose-stimulated insulin release is enhanced, both *in vivo* and *ex vivo* in isolated islets.

### 4.1 Maintained Ca^2+^ dynamics and ER function are essential for adipocyte hormone secretion

The ER is the first compartment of the secretory pathway, where proteins are folded, oligomerized and finally packaged into vesicles that continue their route towards the plasma membrane. Protein folding and oligomerization in ER is controlled by Ca^2+^- dependent chaperones [3] and obesity-associated ER stress has been shown to impair adipocyte hormone secretion [1]. In agreement with this, we found that circulating levels of adiponectin are reduced in the adipocyte-specific SERCA2 null mice (**Fig. 5A**). However, the HMW form of adiponectin remained unaffected in serum from SERCA2 ablated mice (**Fig. 5C**). A few studies suggest that reduced circulating levels of specifically the HMW form of adiponectin is connected to development of obesity-associated metabolic disease and that higher-order adiponectin complexes protect from progress to T2D [60; 61]. Thus, the adiponectin aberrations observed in the adipocyte-specific SERCA2 null mice do not fully concur with adiponectin alterations reported in metabolic disease. The found increase in *Erp44* and *Ero1-Lα*, two ER chaperones belonging to the protein-disulfide isomerase family may underly this difference [62]. Disulfide bonds form and stabilize the higher-order adiponectin complexes [63] and *Erp44* and *Ero1-Lα* are critically important for the proper ER-located modifications and the secretion of adiponectin. Cyclohexamide-induced inhibition of protein synthesis, which leads to retention of proteins within the ER, resulted in production of more HMW adiponectin, at the expense of smaller adiponectin forms [64]. Thus, we suggest that disturbed Ca^2+^ dynamics and ER stress in SERCA2-ablated adipocytes result in retention of adiponectin within the ER. This, together with the concomitant upregulated expression of *Erp44* and *Ero1-Lα*, promote the assembly of HMW adiponectin at the expense of smaller adiponectin forms. Furthermore, circulating adiponectin levels may not only reflect the secretory function of individual adipocytes, but rather the sum of the functionality of different fat depots (as well as effects of negative feedback and hepatic clearance). GWAT and IWAT display different adipocyte hormone release patterns [65; 66] and in obese subjects, small sized visceral adipocytes were positively correlated to serum HMW adiponectin [67]. The finding that SERCA2 knockout GWAT adipocytes displayed a dramatic increase in *Serca3* expression (**Fig. 3B**) points to a less pronounced phenotype in this fat depot. Thus, the higher HMW/total adiponectin in adipocyte-specific SERCA2 knockout mice, may result also from GWAT adipocytes remaining more functional and contributing more to the levels of higher-order adiponectin complexes.

The circulating levels of resistin are dramatically reduced in the adipocyte-specific SERCA2 knockouts (**Fig. 5D**). This is not suprising as resistin, like adiponectin, circulates in differently sized molecular forms [68] and maintained ER chaperone activity is likely essential for its assembly and secretion. Moreover, it has been suggested that resistin itself functions as a chaperone and is retained within the ER during ER stress to protect from cell apoptosis [69]. Leptin, on the other hand, is released from adipocytes as molecular monomers and yet circulating leptin levels are also lower in adipocyte-specific SERCA2 knockouts. We attribute this difference in leptin levels primarily to the reduced adipose tissue mass in the adipocyte-specific SERCA2 knockouts [70]. In support of this, serum leptin levels were similar in HFD-fed wild type and adipocyte-specific SERCA2 knockout mice, the latter with increased adipocyte size which is strongly associated with increased leptin secretion and levels [71; 72]. Thus, our study indicate that loss of adipocyte SERCA2 in obesity/T2D can mediate alterations in adipokine levels although genetic SERCA2 ablation causes more dramatic effects than those observed in metabolic disease where the reduction in adipocyte SERCA2 is less dramatic (**Fig 1E, F, J**).

### 4.2 Genetic ablation of SERCA2 impairs calcium homeostasis and induces different thermogenic adaptations in white and brown adipocytes

Adipocyte-specific SERCA2 knockout BAT and IWAT both display reduced expression of *Prdm16* and *Dio2* (**Suppl. Fig. 6A-B**) and increased mitochondrial UCP1 levels (**Fig. 6F, J**). However, BAT glucose uptake and mitochondrial respiration are reduced while we observed accelerated metabolism in IWAT of the adipocyte-specific SERCA2 knockout mice (**Fig. 6A, G, H, K)**. These differences in mitochondrial respiration between genotype are seen also in cultured white and brown adipocytes (**Fig. 6I, L**) arguing for a cell autonomous mechanism. Similar differences in IWAT and BAT mitochondrial responses to stressors have been reported previously [26; 73; 74]. Moreover, the OCR was recently reported to increase in subcutaneous adipocytes from insulin resistant compared to insulin sensitive obese subjects [75], which is consistent with our findings (**Fig. 1J-K, 6G-H**). In contrast to genetic SERCA2 ablation, acute pharmacological SERCA inhibition in 3T3-L1 adipocytes increases the mitochondrial Ca^2+^ levels associated with increased ROS production and reduced OCR (**Fig. 6B-D**). Thus the browning-like effect in SERCA2-deficient adipocytes appears to be part of an adaptive ER stress-induced mitohormetic that does not occur upon acute SERCA inhibition. Based on several reports [76–80], we propose that the upregulation of UCP1 (**Fig. 6F, J**) is triggered by increased mitochondrial ROS levels, in order to suppress further increases in the ROS production and Ca^2+^ overload in SERCA2- deficient adipocytes. It is also possible that the already high levels of UCP1 and endogenous antioxidant enzymes prevent the mitohormetic response in BAT, as has been suggested previously [74]. The different adaptations of white and brown adipocytes to SERCA2 ablation may also be explained by differential induction of FGF21; *Fgf21* expression was increased in IWAT but not in BAT (**Fig. 6N**). However, the reduction in mitochondrial metabolism may also signpost a specific role of SERCA2 in mitochondrial function of brown adipocytes that cannot be compensated for by SERCA1 and 3.

SERCA2 has been implicated in ATP-dependent thermogenesis by Ca^2+^ cycling that can compensate for loss of UCP1 in beige, but not brown, adipose tissue [81]. The finding that the norepinephrine-induced increase in OCR was normal in SERCA2- deficient brown adipocytes (**Suppl. Fig. 6C**) are in line with that UCP1 is essential for norepinephrine-induced thermogenesis in BAT [82]. It is however possible that adrenergically stimulated thermogenesis is slightly blunted in SERCA2-deficient white adipocytes. Alternatively, the upregulation of UCP1 that we observe (**Fig. 6F**) serves to compensate for the lack of this thermogenic mechanism in SERCA2-deficient adipocytes.

### 4.3 SERCA2 in fat accumulation

The lipodystrophic phenotype of adipocyte-specific SERCA2 knockout mice is in agreement with the proposed role of SERCA and Ca^2+^ homesostasis in fat storage [83]. However, our data suggest that fat storage and the capacity for *de novo* lipogenesis is normal in SERCA2 deficient adipocytes (**Fig. 6A**). This proposes that SERCA2 is not essential for fat storage in mouse adipocytes, and that loss of SERCA2 rather makes the white adipocytes less viable, as supported by the increased crown-like structures and the trend towards increased *Casp3* levels under chow conditions (**Fig. 3F, 4M**) and fewer, but larger, adipocytes under HFD-fed obese conditions (**Fig. 4K-L**). Thus, both increased apoptosis and accelerated metabolism (as discussed above) provide plausible explanations for the reduced fat mass in these mice.

### 4.4 Conclusion

Collectively, our data emphasizes the importance of SERCA2 for preserved calcium homostasis associated with proper ER and mitochondrial function in white and brown adipocytes. In obesity, reduction of adipocyte SERCA2 may thus contribute to adipose tissue dysfunction and the pathogenesis of metabolic disorders.

## Acknowledgments

This study was funded by the Swedish Research Council (2013-07107, 2017-00792, 2019-01239 and 2020-01463), the Magnus Bergvall foundation (2016-01711), the NovoNordisk Foundation (NNF19OC0056601), and the Swedish Diabetes Foundation (DIA2016-127, DIA2018-358, DIA2019-419, DIA2014-074, DIA2015-062, DIA2017-273 and DIA2018-354). The SERCA2^flox^ mouse model was a kind gift from Geir Christensen and William Louch (Institute for Experimental Medical Research at Oslo University Hospital Ullevål, Oslo, Norway). Graphical abstract was created with BioRender.com.

## Author contributions

IWA and CO conceived the idea, supervised this work, interpreted data and wrote the manuscript. They have contributed equally to this study and are shared senior authors. MBT and EB performed experiments and interpreted data, made figures and wrote parts of the manuscript. BC, EP, YW, SM and CJ performed experiments and assisted in data interpretation. CJ an PS provided human adipocyte samples and clinical data. PR and PS interpreted data and gave valuable feedback to this work. All authors have approved the final version of this manuscript.

---

**Supplemental figure 1.**
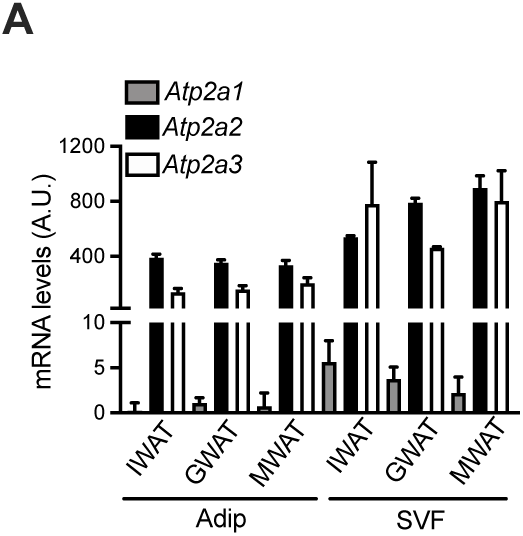
SERCA2 and 3 are the predominant SERCA paralogs in the adipose tissue. *Atp2a1* (*Serca1)*, *Atp2a2* (Serca*2)* and *Atp2a3* (Serca*3)* gene expression in the adipocyte and the stromal vascular fractions (SVF) of inguinal (IWAT), gonadal (GWAT) and mesenteric white adipose tissue (MWAT) from chow-fed mice (N=4-6).

**Supplemental figure 2.**
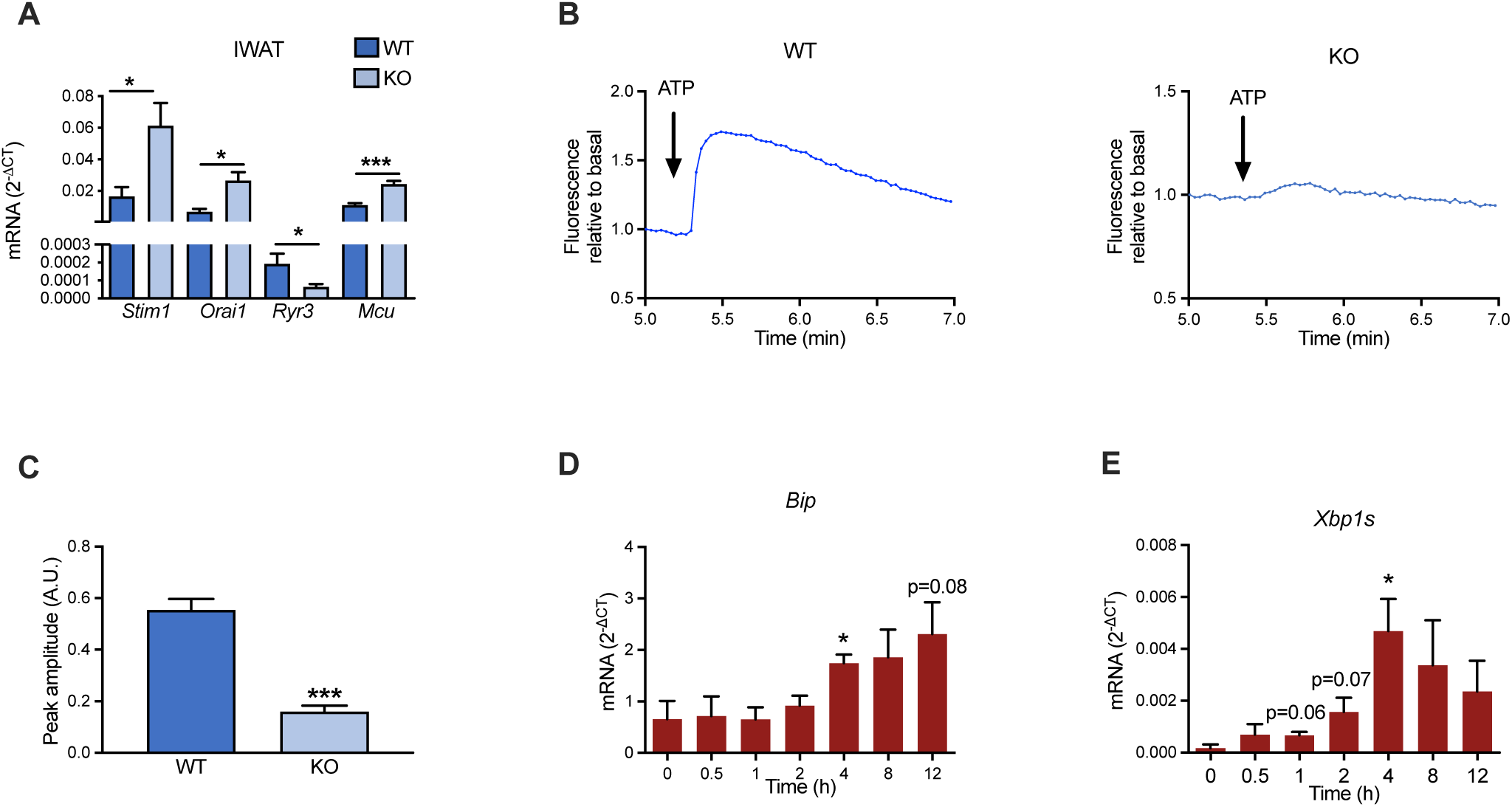
Inhibition of SERCA in adipocytes is associated with altered Ca^2+^ homeostasis and ER-stress. (**A**) Calcium transporter gene expression markers in IWAT in chow-fed Wildtype (WT) and adipocyte specific SERCA2 Knockout (KO) mice (N=6-10). (**B**) Example traces of mitochondrial Ca^2+^ and (**C)** ATP peak amplitude of the mitochondrial Ca^2+^ in response to ATP in differentiated WT and adipocyte-specific SERCA2 KO IWAT SVF adipocytes. *Bip* (**D**) and *Xbp1s* (**E**) mRNA levels in 3T3-L1 adipocytes in response to 20 μM CPA (*p<0.05 vs. time 0h). All values (N=3-6) are expressed as mean ± SEM; *p<0.05, **p<0.01, ***p<0.001.

**Supplemental figure 3.**
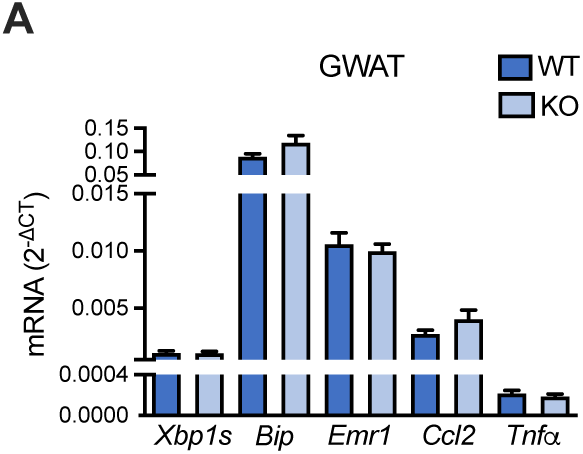
ER stress and inflammation markers are not altered in GWAT from adipocyte- specifi SERCA2 knockout mice. ER stress and inflammation gene expression markers in GWAT in chow-fed WT and adipocyte-specifi SERCA2 KO mice (N=6-10). All values are expressed as mean ± SEM; *p<0.05, **p<0.01, ***p<0.001 vs WT.

**Supplemental figure 4.**
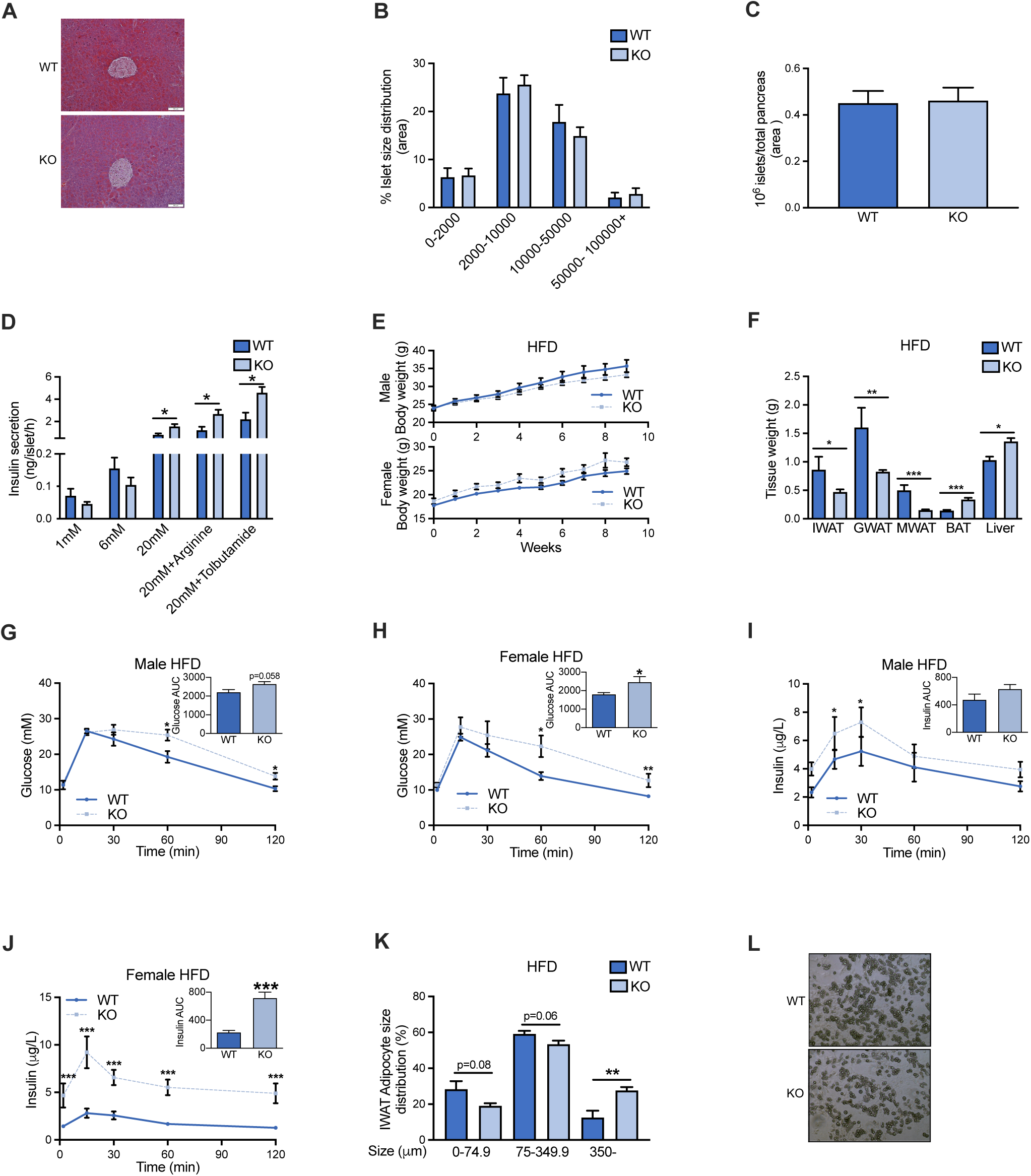
Adipocyte-specifi SERCA2 knockout mice are glucose intolerant, but display increased glucose-stimulated insulin release. (**A**) Representative pancreas H&E sections, (**B**) % islets size distribution and (**C**) number of islets in pancreas sections from chow-fed WT and adipocyte-specifi SERCA2 KO mice. (**D**) Insulin secretion from pancreatic islets isolated from chow-fed WT and adipocyte-specifi SERCA2 KO mice (n= 3 experiments; n= 4 mice per genotype) in the presence of different concentration of glucose and Arginine or Tolbutamide, as indicated. (**E**) Body weight of male and female of WT and adipocyte-specifi SERCA2 KO mice during 10-week high fat diet (HFD) time course. Male and female (**F**) IWAT, GWAT, MWAT, BAT and liver weight, (**G-H**) glucose and (**I-J**) insulin levels in response to an oral glucose load, and (**K**) IWAT adipocyte size distribution in 10-week-HFD-fed WT and adipocyte-specifi SERCA2 KO mice. (**L**) Representative picture at day 8 of adipocytes differentiated from isolated WT and adipocyte-specifi SERCA2 KO IWAT SVF. All values (N=6-10) are expressed as mean ± SEM *p<0.05, **p<0.01, ***p<0.001 for WT vs KO.

**Supplemental figure 5.**
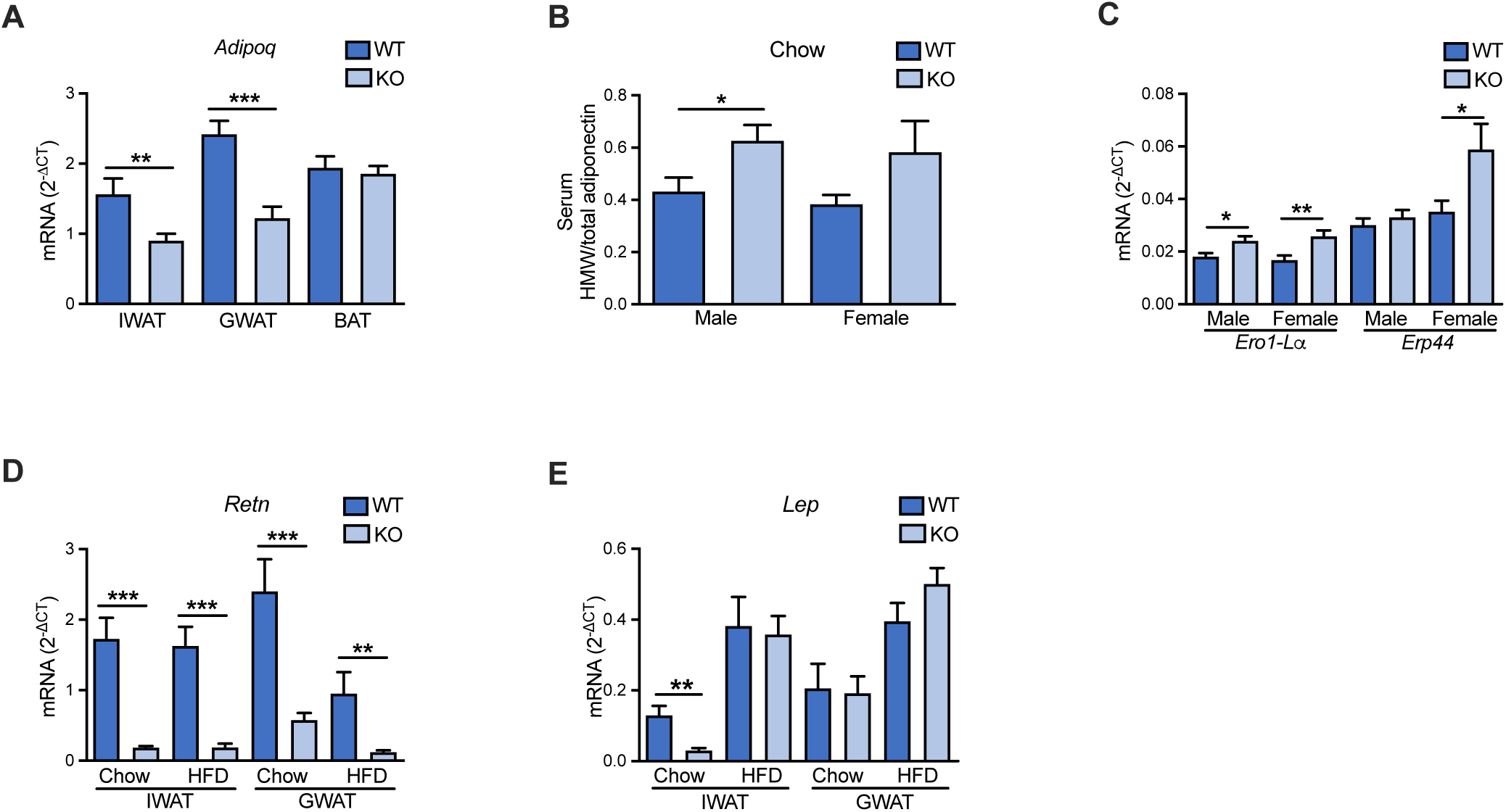
ER stress markers and adipocyte hormones are altered in adipocyte-specifi SERCA2 knockout mice. (**A**) Adiponectin (*Adipoq*) expression in IWAT, GWAT and BAT in 10-week-HFD-fed WT and adipocyte-specifi SERCA2 KO mice. (**B**) Serum HMW-Total adiponectin ratio in male and female chow- fed WT and adipocyte-specifi SERCA2 KO mice. (**C**) *Ero1-Lα* and *Erp44* gene expression in male and female chow-fed WT and adipocyte-specifi SERCA2 KO mice. IWAT and GWAT (**D**) Resistin (*Retn*) and (**E**) leptin (*Lep*) gene expression in chow and 10-week-HFD-fed WT and adipocyte-specifi SERCA2 KO mice. All values (N=6-10) are expressed as mean ± SEM *p<0.05, **p<0.01, ***p<0.001 for WT vs KO.

**Supplemental figure 6.**
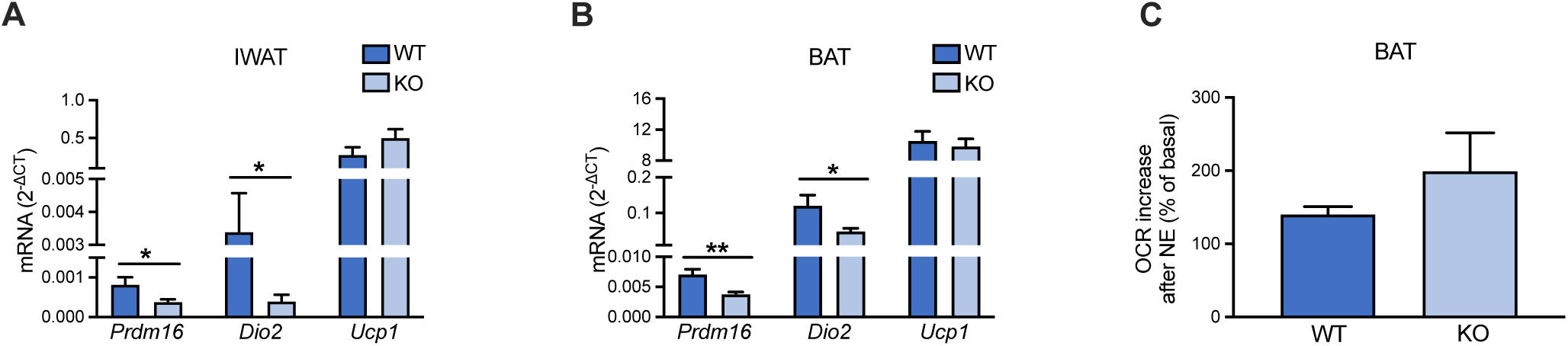
Effect of adipocyte-specifi SERCA2 ablation on adipose tissue browning markers in white and brown adipose tissue. *Prdm16*, *Dio2* and *Ucp1* expression in (**A**) IWAT and (**B**) BAT in chow-fed WT and adipocyte-specifi SERCA2 KO mice. (**C**) Norepinephrine (NE, 1µM)-induced increase in oxygen consumption rate (OCR) in brown adipocytes differentiated from BAT SVF of chow-fed WT and adipocyte- specifi SERCA2 KO mice. All values (N=3-6) are expressed as mean ± SEM *p<0.05, **p<0.01, ***p<0.001 for WT vs KO.

**Supplementary Table 1.**
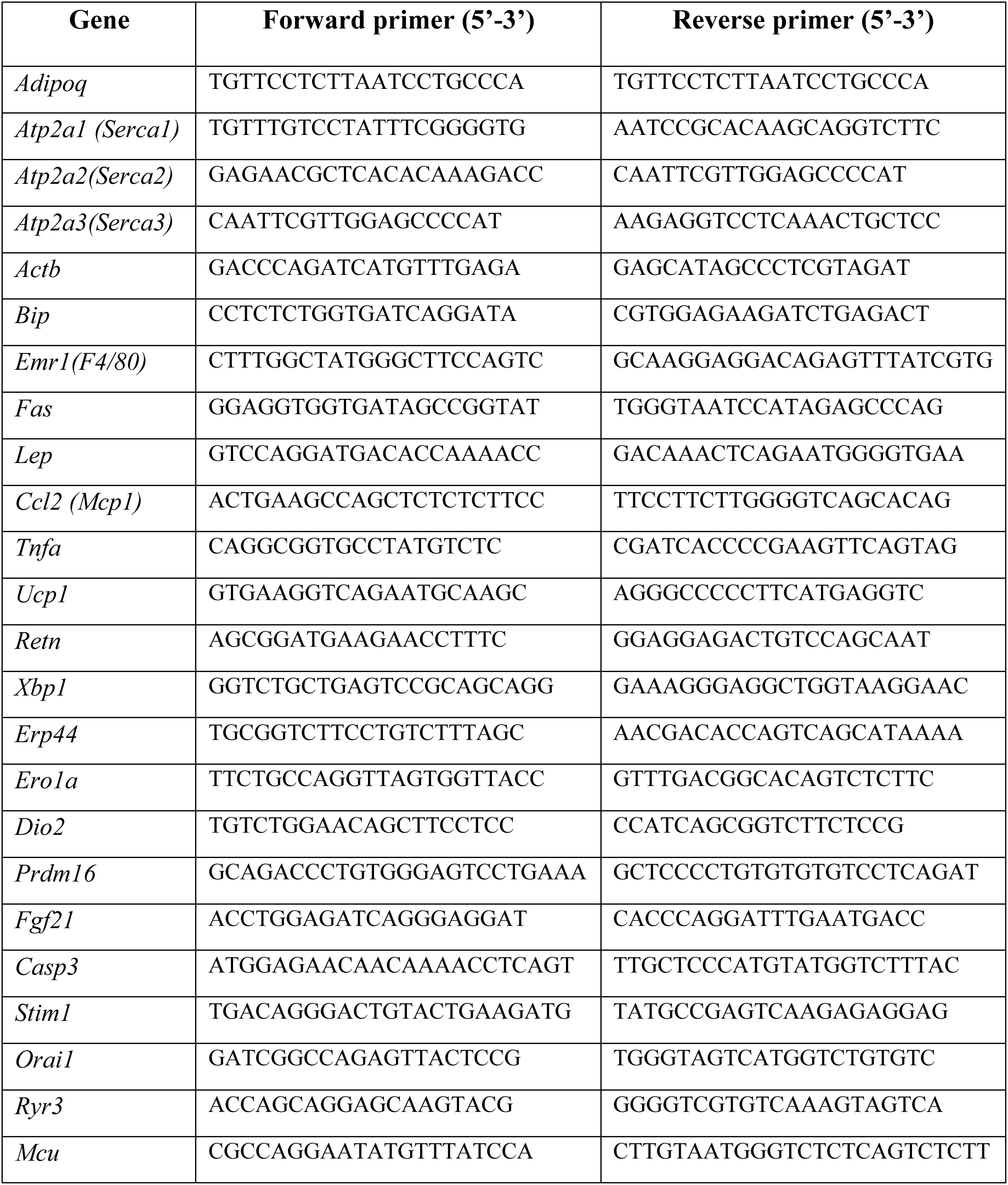

## References

1. Longo, M., Zatterale, F., Naderi, J., Parrillo, L., Formisano, P., Raciti, G.A., et al., 2019. Adipose Tissue Dysfunction as Determinant of Obesity-Associated Metabolic Complications. International Journal of Molecular Sciences 20(9):2358 DOI: 10.3390/ijms20092358.

2. Liang, W., Ye, D.d., 2019. The potential of adipokines as biomarkers and therapeutic agents for vascular complications in type 2 diabetes mellitus. Cytokine & Growth Factor Reviews 48:32–39 DOI: https://doi.org/10.1016/j.cytogfr.2019.06.002.

3. Hetz, C., Zhang, K., Kaufman, R.J., 2020. Mechanisms, regulation and functions of the unfolded protein response. Nature reviews. Molecular cell biology 21(8):421–438 DOI: 10.1038/s41580-020-0250-z.

4. Izawa, T., Komabayashi, T., 1994. Ca2+ and lipolysis in adipocytes from exercise-trained rats. J Appl Physiol (1985) 77(6):2618–2624.

5. Ohisalo, J.J., 1980. Modulation of lipolysis by adenosine and Ca2+ in fat cells from hypothyroid rats. FEBS Lett 116(1):91–94.

6. Fedorenko, O.A., Pulbutr, P., Banke, E., Akaniro-Ejim, N.E., Bentley, D.C., Olofsson, C.S., et al., 2020. CaV1.2 and CaV1.3 voltage-gated L-type Ca2+ channels in rat white fat adipocytes. J Endocrinol 244(2):369–381 DOI: 10.1530/JOE-19-0493.

7. Worrall, D.S., Olefsky, J.M., 2002. The effects of intracellular calcium depletion on insulin signaling in 3T3-L1 adipocytes. Mol Endocrinol 16(2):378–389 DOI: 10.1210/mend.16.2.0776.

8. Draznin, B., 1993. Cytosolic Calcium and Insulin Resistance. American Journal of Kidney Diseases 21(6, Supplement):S32–S38 DOI: https://doi.org/10.1016/0272-6386(93)70122-F.

9. Wang, Y., Ali, Y., Lim, C.Y., Hong, W., Pang, Z.P., Han, W., 2014. Insulin-stimulated leptin secretion requires calcium and PI3K/Akt activation. Biochem J 458(3):491–498 DOI: 10.1042/BJ20131176.

10. Cammisotto, P.G., Bukowiecki, L.J., 2004. Role of calcium in the secretion of leptin from white adipocytes. Am J Physiol Regul Integr Comp Physiol 287(6):R1380–1386.

11. Levy, J.R., Gyarmati, J., Lesko, J.M., Adler, R.A., Stevens, W., 2000. Dual regulation of leptin secretion: intracellular energy and calcium dependence of regulated pathway. Am J Physiol Endocrinol Metab 278(5):E892–901.

12. El Hachmane, M.F., Komai, A.M., Olofsson, C.S., 2015. Cooling Reduces cAMP-Stimulated Exocytosis and Adiponectin Secretion at a Ca2+-Dependent Step in 3T3-L1 Adipocytes. PLoS One 10(3):e0119530 DOI: 10.1371/journal.pone.0119530.

13. Komai, A.M., Brannmark, C., Musovic, S., Olofsson, C.S., 2014. PKA-independent cAMP stimulation of white adipocyte exocytosis and adipokine secretion: modulations by Ca2+ and ATP. J Physiol 592(Pt 23):5169–5186 DOI: 10.1113/jphysiol.2014.280388.

14. Waldeck-Weiermair, M., Deak, A.T., Groschner, L.N., Alam, M.R., Jean-Quartier, C., Malli, R., et al., 2013. Molecularly distinct routes of mitochondrial Ca2+ uptake are activated depending on the activity of the sarco/endoplasmic reticulum Ca2+ ATPase (SERCA). J Biol Chem 288(21):15367–15379 DOI: 10.1074/jbc.M113.462259.

15. Arruda, A.P., Hotamisligil, G.S., 2015. Calcium Homeostasis and Organelle Function in the Pathogenesis of Obesity and Diabetes. Cell Metab 22(3):381–397 DOI: 10.1016/j.cmet.2015.06.010.

16. Rizzuto, R., De Stefani, D., Raffaello, A., Mammucari, C., 2012. Mitochondria as sensors and regulators of calcium signalling. Nature Reviews Molecular Cell Biology 13(9):566–578 DOI: 10.1038/nrm3412.

17. Fu, S., Yang, L., Li, P., Hofmann, O., Dicker, L., Hide, W., et al., 2011. Aberrant lipid metabolism disrupts calcium homeostasis causing liver endoplasmic reticulum stress in obesity. Nature 473(7348):528–531 DOI: 10.1038/nature09968.

18. Park, S.W., Zhou, Y., Lee, J., Lee, J., Ozcan, U., 2010. Sarco(endo)plasmic reticulum Ca2+-ATPase 2b is a major regulator of endoplasmic reticulum stress and glucose homeostasis in obesity. Proc Natl Acad Sci U S A 107(45):19320–19325 DOI: 10.1073/pnas.1012044107.

19. Varadi, A., Molnar, E., Ostenson, C.G., Ashcroft, S.J., 1996. Isoforms of endoplasmic reticulum Ca(2+)-ATPase are differentially expressed in normal and diabetic islets of Langerhans. Biochem J 319 ( Pt 2):521–527.

20. Waller, A.P., Kalyanasundaram, A., Hayes, S., Periasamy, M., Lacombe, V.A., 2015. Sarcoplasmic reticulum Ca2+ ATPase pump is a major regulator of glucose transport in the healthy and diabetic heart. Biochimica et Biophysica Acta (BBA) - Molecular Basis of Disease 1852(5):873–881 DOI: https://doi.org/10.1016/j.bbadis.2015.01.009.

21. Kang, S., Dahl, R., Hsieh, W., Shin, A., Zsebo, K.M., Buettner, C., et al., 2016. Small Molecular Allosteric Activator of the Sarco/Endoplasmic Reticulum Ca2+- ATPase (SERCA) Attenuates Diabetes and Metabolic Disorders. J Biol Chem 291(10):5185–5198 DOI: 10.1074/jbc.M115.705012.

22. Honnor, R.C., Dhillon, G.S., Londos, C., 1985. cAMP-dependent protein kinase and lipolysis in rat adipocytes. I. Cell preparation, manipulation, and predictability in behavior. J Biol Chem 260(28):15122–15129.

23. Bauzá-Thorbrügge, M., M. Galmés-Pascual, B., Sbert-Roig, M., J. García-Palmer, F., Gianotti, M., M. Proenza, A., et al., 2017. Antioxidant peroxiredoxin 3 expression is regulated by 17beta-estradiol in rat white adipose tissue. The Journal of Steroid Biochemistry and Molecular Biology 172:9–19 DOI: https://doi.org/10.1016/j.jsbmb.2017.05.008.

24. Jonsson, C., Castor Batista, A.P., Kjolhede, P., Stralfors, P., 2019. Insulin and beta-adrenergic receptors mediate lipolytic and anti-lipolytic signalling that is not altered by type 2 diabetes in human adipocytes. Biochem J 476(19):2883–2908 DOI: 10.1042/BCJ20190594.

25. Stralfors, P., Honnor, R.C., 1989. Insulin-induced dephosphorylation of hormone-sensitive lipase. Correlation with lipolysis and cAMP-dependent protein kinase activity. Eur J Biochem 182(2):379–385.

26. Peris, E., Micallef, P., Paul, A., Palsdottir, V., Enejder, A., Bauza-Thorbrugge, M., et al., 2019. Antioxidant treatment induces reductive stress associated with mitochondrial dysfunction in adipocytes. J Biol Chem 294(7):2340–2352 DOI: 10.1074/jbc.RA118.004253.

27. Rogers, G.W., Brand, M.D., Petrosyan, S., Ashok, D., Elorza, A.A., Ferrick, D.A., et al., 2011. High throughput microplate respiratory measurements using minimal quantities of isolated mitochondria. PloS one 6(7):e21746–e21746 DOI: 10.1371/journal.pone.0021746.

28. Astrom-Olsson, K., Li, L., Olofsson, C.S., Boren, J., Ohlin, H., Grip, L., 2012. Impact of hypoxia, simulated ischemia and reperfusion in HL-1 cells on the expression of FKBP12/FKBP12.6 and intracellular calcium dynamics. Biochem Biophys Res Commun 422(4):732–738 DOI: S0006-291X(12)00948-5 [pii] 10.1016/j.bbrc.2012.05.071.

29. Grynkiewicz, G., Poenie, M., Tsien, R.Y., 1985. A new generation of Ca2+ indicators with greatly improved fluorescence properties. J Biol Chem 260(6):3440–3450.

30. Maxwell, J.T., Tsai, C.-H., Mohiuddin, T.A., Kwong, J.Q., 2018. Analyses of Mitochondrial Calcium Influx in Isolated Mitochondria and Cultured Cells. Journal of visualized experiments : JoVE(134):57225 DOI: 10.3791/57225.

31. Bidault, G., Garcia, M., Vantyghem, M.C., Ducluzeau, P.H., Morichon, R., Thiyagarajah, K., et al., 2013. Lipodystrophy-linked LMNA p.R482W mutation induces clinical early atherosclerosis and in vitro endothelial dysfunction. Arterioscler Thromb Vasc Biol 33(9):2162–2171 DOI: 10.1161/atvbaha.113.301933.

32. Kulkarni, R.N., Roper, M.G., Dahlgren, G., Shih, D.Q., Kauri, L.M., Peters, J.L., et al., 2004. Islet secretory defect in insulin receptor substrate 1 null mice is linked with reduced calcium signaling and expression of sarco(endo)plasmic reticulum Ca2+-ATPase (SERCA)-2b and -3. Diabetes 53(6):1517–1525.

33. Wold, L.E., Dutta, K., Mason, M.M., Ren, J., Cala, S.E., Schwanke, M.L., et al., 2005. Impaired SERCA function contributes to cardiomyocyte dysfunction in insulin resistant rats. J Mol Cell Cardiol 39(2):297–307 DOI: 10.1016/j.yjmcc.2005.03.014.

34. Rabinovich-Nikitin, I., Kirshenbaum, L.A., 2019. Hypoxia-inducible factor 1 regulates SERCA2 in the heart by modulating miR-29c levels. Am J Physiol Heart Circ Physiol 316(5):H1211–H1213 DOI: 10.1152/ajpheart.00019.2019.

35. Revuelta-Lopez, E., Cal, R., Herraiz-Martinez, A., de Gonzalo-Calvo, D., Nasarre, L., Roura, S., et al., 2015. Hypoxia-driven sarcoplasmic/endoplasmic reticulum calcium ATPase 2 (SERCA2) downregulation depends on low-density lipoprotein receptor-related protein 1 (LRP1)-signalling in cardiomyocytes. J Mol Cell Cardiol 85:25–36 DOI: 10.1016/j.yjmcc.2015.04.028.

36. Ronkainen, V.P., Skoumal, R., Tavi, P., 2011. Hypoxia and HIF-1 suppress SERCA2a expression in embryonic cardiac myocytes through two interdependent hypoxia response elements. J Mol Cell Cardiol 50(6):1008–1016 DOI: 10.1016/j.yjmcc.2011.02.017.

37. Hosogai, N., Fukuhara, A., Oshima, K., Miyata, Y., Tanaka, S., Segawa, K., et al., 2007. Adipose tissue hypoxia in obesity and its impact on adipocytokine dysregulation. Diabetes 56(4):901–911 DOI: 10.2337/db06-0911.

38. Wang, B., Wood, I.S., Trayhurn, P., 2007. Dysregulation of the expression and secretion of inflammation-related adipokines by hypoxia in human adipocytes. Pflugers Arch 455(3):479–492 DOI: 10.1007/s00424-007-0301-8.

39. Várnai, P., Hunyady, L., Balla, T., 2009. STIM and Orai: the long-awaited constituents of store-operated calcium entry. Trends Pharmacol Sci 30(3):118–128 DOI: 10.1016/j.tips.2008.11.005.

40. Laver, D.R., 2007. Ca2+ stores regulate ryanodine receptor Ca2+ release channels via luminal and cytosolic Ca2+ sites. Biophysical journal 92(10):3541–3555 DOI: 10.1529/biophysj.106.099028.

41. El Hachmane, M.F., Ermund, A., Brannmark, C., Olofsson, C.S., 2018. Extracellular ATP activates store-operated Ca(2+) entry in white adipocytes: functional evidence for STIM1 and ORAI1. Biochem J 475(3):691–704 DOI: 10.1042/BCJ20170484.

42. Lee, H., Jun, D.J., Suh, B.C., Choi, B.H., Lee, J.H., Do, M.S., et al., 2005. Dual roles of P2 purinergic receptors in insulin-stimulated leptin production and lipolysis in differentiated rat white adipocytes. J Biol Chem 280(31):28556–28563 DOI: 10.1074/jbc.M411253200.

43. Missiroli, S., Patergnani, S., Caroccia, N., Pedriali, G., Perrone, M., Previati, M., et al., 2018. Mitochondria-associated membranes (MAMs) and inflammation. Cell death & disease 9(3):329–329 DOI: 10.1038/s41419-017-0027-2.

44. Liao, Y., Hao, Y., Chen, H., He, Q., Yuan, Z., Cheng, J., 2015. Mitochondrial calcium uniporter protein MCU is involved in oxidative stress-induced cell death. Protein & cell 6(6):434–442 DOI: 10.1007/s13238-015-0144-6.

45. Komai, A.M., Musovic, S., Peris, E., Alrifaiy, A., El Hachmane, M.F., Johansson, M., et al., 2016. White Adipocyte Adiponectin Exocytosis Is Stimulated via beta3-Adrenergic Signaling and Activation of Epac1: Catecholamine Resistance in Obesity and Type 2 Diabetes. Diabetes 65(11):3301–3313 DOI: 10.2337/db15-1597.

46. Koh, E.H., Park, J.Y., Park, H.S., Jeon, M.J., Ryu, J.W., Kim, M., et al., 2007. Essential role of mitochondrial function in adiponectin synthesis in adipocytes. Diabetes 56(12):2973–2981 DOI: 10.2337/db07-0510.

47. Zhou, L., Liu, M., Zhang, J., Chen, H., Dong, L.Q., Liu, F., 2010. DsbA-L alleviates endoplasmic reticulum stress-induced adiponectin downregulation. Diabetes 59(11):2809–2816 DOI: 10.2337/db10-0412.

48. Wang, Q.A., Tao, C., Gupta, R.K., Scherer, P.E., 2013. Tracking adipogenesis during white adipose tissue development, expansion and regeneration. Nat Med 19(10):1338–1344 DOI: 10.1038/nm.3324.

49. Qiang, L., Wang, H., Farmer, S.R., 2007. Adiponectin secretion is regulated by SIRT1 and the endoplasmic reticulum oxidoreductase Ero1-L alpha. Mol Cell Biol 27(13):4698–4707 DOI: 10.1128/MCB.02279-06.

50. Wang, Z.V., Scherer, P.E., 2008. DsbA-L is a versatile player in adiponectin secretion. Proc Natl Acad Sci U S A 105(47):18077–18078 DOI: 10.1073/pnas.0810027105.

51. Tripathi, D., Kant, S., Pandey, S., Ehtesham, N.Z., 2020. Resistin in metabolism, inflammation, and disease. The FEBS Journal 287(15):3141–3149 DOI: https://doi.org/10.1111/febs.15322.

52. Steppan, C.M., Bailey, S.T., Bhat, S., Brown, E.J., Banerjee, R.R., Wright, C.M., et al., 2001. The hormone resistin links obesity to diabetes. Nature 409(6818):307–312 DOI: 10.1038/35053000.

53. Asterholm, I.W., Rutkowski, J.M., Fujikawa, T., Cho, Y.R., Fukuda, M., Tao, C., et al., 2014. Elevated resistin levels induce central leptin resistance and increased atherosclerotic progression in mice. Diabetologia 57(6):1209–1218 DOI: 10.1007/s00125-014-3210-3.

54. Patel, S.D., Rajala, M.W., Rossetti, L., Scherer, P.E., Shapiro, L., 2004. Disulfide-Dependent Multimeric Assembly of Resistin Family Hormones. Science 304(5674):1154–1158 DOI: 10.1126/science.1093466.

55. Musovic, S., Shrestha, M.M., Komai, A.M., Olofsson, C.S., 2021. Resistin is co-secreted with adiponectin in white mouse adipocytes. Biochem Biophys Res Commun 534:707–713 DOI: https://doi.org/10.1016/j.bbrc.2020.11.013.

56. Kim, S.H., Kim, K.H., Kim, H.-K., Kim, M.-J., Back, S.H., Konishi, M., et al., 2015. Fibroblast growth factor 21 participates in adaptation to endoplasmic reticulum stress and attenuates obesity-induced hepatic metabolic stress. Diabetologia 58(4):809–818 DOI: 10.1007/s00125-014-3475-6.

57. Schaap, F.G., Kremer, A.E., Lamers, W.H., Jansen, P.L.M., Gaemers, I.C., 2013. Fibroblast growth factor 21 is induced by endoplasmic reticulum stress. Biochimie 95(4):692–699 DOI: https://doi.org/10.1016/j.biochi.2012.10.019.

58. Fisher, F.M., Kleiner, S., Douris, N., Fox, E.C., Mepani, R.J., Verdeguer, F., et al., 2012. FGF21 regulates PGC-1α and browning of white adipose tissues in adaptive thermogenesis. Genes Dev 26(3):271–281 DOI: 10.1101/gad.177857.111.

59. Lee, P., Linderman, Joyce D., Smith, S., Brychta, Robert J., Wang, J., Idelson, C., et al., 2014. Irisin and FGF21 Are Cold-Induced Endocrine Activators of Brown Fat Function in Humans. Cell Metab 19(2):302–309 DOI: https://doi.org/10.1016/j.cmet.2013.12.017.

60. Goto, M., Goto, A., Morita, A., Deura, K., Sasaki, S., Aiba, N., et al., 2014. Low-molecular-weight adiponectin and high-molecular-weight adiponectin levels in relation to diabetes. Obesity (Silver Spring) 22(2):401–407 DOI: 10.1002/oby.20553.

61. Seino, Y., Hirose, H., Saito, I., Itoh, H., 2009. High-molecular-weight adiponectin is a predictor of progression to metabolic syndrome: a population-based 6-year follow-up study in Japanese men. Metabolism 58(3):355–360 DOI: https://doi.org/10.1016/j.metabol.2008.10.008.

62. Hampe, L., Radjainia, M., Xu, C., Harris, P.W.R., Bashiri, G., Goldstone, D.C., et al., 2015. Regulation and Quality Control of Adiponectin Assembly by Endoplasmic Reticulum Chaperone ERp44. J Biol Chem 290(29):18111–18123 DOI: 10.1074/jbc.M115.663088.

63. Tsao, T.S., Tomas, E., Murrey, H.E., Hug, C., Lee, D.H., Ruderman, N.B., et al., 2003. Role of disulfide bonds in Acrp30/adiponectin structure and signaling specificity. Different oligomers activate different signal transduction pathways. J Biol Chem 278(50):50810–50817 DOI: 10.1074/jbc.M309469200.

64. Wang, Z.V., Schraw, T.D., Kim, J.Y., Khan, T., Rajala, M.W., Follenzi, A., et al., 2007. Secretion of the adipocyte-specific secretory protein adiponectin critically depends on thiol-mediated protein retention. Mol Cell Biol 27(10):3716–3731 DOI: 10.1128/MCB.00931-06.

65. Scheja, L., Heeren, J., 2019. The endocrine function of adipose tissues in health and cardiometabolic disease. Nature Reviews Endocrinology 15(9):507–524 DOI: 10.1038/s41574-019-0230-6.

66. Meyer, L.K., Ciaraldi, T.P., Henry, R.R., Wittgrove, A.C., Phillips, S.A., 2013. Adipose tissue depot and cell size dependency of adiponectin synthesis and secretion in human obesity. Adipocyte 2(4):217–226 DOI: 10.4161/adip.24953.

67. Reneau, J., Goldblatt, M., Gould, J., Kindel, T., Kastenmeier, A., Higgins, R., et al., 2018. Effect of adiposity on tissue-specific adiponectin secretion. PLoS One 13(6):e0198889 DOI: 10.1371/journal.pone.0198889.

68. Chen, J., Wang, L., Boeg, Y.S., Xia, B., Wang, J., 2002. Differential dimerization and association among resistin family proteins with implications for functional specificity. Journal of Endocrinology 175(2):499–504 DOI: 10.1677/joe.0.1750499.

69. Suragani, M., Aadinarayana, V.D., Pinjari, A.B., Tanneeru, K., Guruprasad, L., Banerjee, S., et al., 2013. Human resistin, a proinflammatory cytokine, shows chaperone-like activity. Proc Natl Acad Sci U S A 110(51):20467–20472 DOI: 10.1073/pnas.1306145110.

70. Frederich, R.C., Hamann, A., Anderson, S., Löllmann, B., Lowell, B.B., Flier, J.S., 1995. Leptin levels reflect body lipid content in mice: Evidence for diet-induced resistance to leptin action. Nat Med 1(12):1311–1314 DOI: 10.1038/nm1295-1311.

71. Zhang, Y., Guo, K.-Y., Diaz, P.A., Heo, M., Leibel, R.L., 2002. Determinants of leptin gene expression in fat depots of lean mice. American Journal of Physiology-Regulatory, Integrative and Comparative Physiology 282(1):R226–R234 DOI: 10.1152/ajpregu.00392.2001.

72. Skurk, T., Alberti-Huber, C., Herder, C., Hauner, H., 2007. Relationship between Adipocyte Size and Adipokine Expression and Secretion. The Journal of Clinical Endocrinology & Metabolism 92(3):1023–1033 DOI: 10.1210/jc.2006-1055.

73. Han, Y.H., Buffolo, M., Pires, K.M., Pei, S., Scherer, P.E., Boudina, S., 2016. Adipocyte-Specific Deletion of Manganese Superoxide Dismutase Protects From Diet-Induced Obesity Through Increased Mitochondrial Uncoupling and Biogenesis. Diabetes 65(9):2639–2651 DOI: 10.2337/db16-0283.

74. Lettieri Barbato, D., Tatulli, G., Aquilano, K., Ciriolo, M.R., 2015. Mitochondrial Hormesis links nutrient restriction to improved metabolism in fat cell. Aging (Albany NY) 7(10):869–881 DOI: 10.18632/aging.100832.

75. Bohm, A., Keuper, M., Meile, T., Zdichavsky, M., Fritsche, A., Haring, H.U., et al., 2020. Increased mitochondrial respiration of adipocytes from metabolically unhealthy obese compared to healthy obese individuals. Sci Rep 10(1):12407 DOI: 10.1038/s41598-020-69016-9.

76. Mailloux, R.J., Harper, M.-E., 2011. Uncoupling proteins and the control of mitochondrial reactive oxygen species production. Free Radical Biology and Medicine 51(6):1106–1115 DOI: https://doi.org/10.1016/j.freeradbiomed.2011.06.022.

77. Lettieri Barbato, D., Tatulli, G., Maria Cannata, S., Bernardini, S., Aquilano, K., Ciriolo, M.R., 2015. Glutathione Decrement Drives Thermogenic Program In Adipose Cells. Scientific reports 5:13091–13091 DOI: 10.1038/srep13091.

78. Oelkrug, R., Kutschke, M., Meyer, C.W., Heldmaier, G., Jastroch, M., 2010. Uncoupling protein 1 decreases superoxide production in brown adipose tissue mitochondria. J Biol Chem 285(29):21961–21968 DOI: 10.1074/jbc.M110.122861.

79. Stier, A., Bize, P., Habold, C., Bouillaud, F., Massemin, S., Criscuolo, F., 2014. Mitochondrial uncoupling prevents cold-induced oxidative stress: a case study using UCP1 knockout mice. Journal of Experimental Biology 217(4):624–630 DOI: 10.1242/jeb.092700.

80. Kazak, L., Chouchani, E.T., Stavrovskaya, I.G., Lu, G.N.Z., Jedrychowski, M.P., Egan, D.F., et al., 2017. UCP1 deficiency causes brown fat respiratory chain depletion and sensitizes mitochondria to calcium overload-induced dysfunction. Proc Natl Acad Sci U S A 114(30):7981–7986 DOI: 10.1073/pnas.1705406114.

81. Ikeda, K., Kang, Q., Yoneshiro, T., Camporez, J.P., Maki, H., Homma, M., et al., 2017. UCP1-independent signaling involving SERCA2b-mediated calcium cycling regulates beige fat thermogenesis and systemic glucose homeostasis. Nat Med 23(12):1454–1465 DOI: 10.1038/nm.4429.

82. Matthias, A., Ohlson, K.B.E., Fredriksson, J.M., Jacobsson, A., Nedergaard, J., Cannon, B., 2000. Thermogenic responses in brown fat cells are fully UCP1- dependent - UCP2 or UCP3 do not substitute for UCP1 in adrenergically or fatty acid-induced thermogenesis. Journal of Biological Chemistry 275(33):25073–25081 DOI: DOI 10.1074/jbc.M000547200.

83. Bi, J., Wang, W., Liu, Z., Huang, X., Jiang, Q., Liu, G., et al., 2014. Seipin Promotes Adipose Tissue Fat Storage through the ER Ca2+-ATPase SERCA. Cell Metabolism 19(5):861–871 DOI: https://doi.org/10.1016/j.cmet.2014.03.028.

